# The ViRA-L (Visual Recognition Assessment–Leuven): Development, Validation, and Age-Stratified Normative Characterization of a Multi-Category Visual Recognition Battery

**DOI:** 10.64898/2026.10.27.761867

**Authors:** Laura Soen, Céline R. Gillebert, Hans Op de Beeck

## Abstract

**Objective:** To develop and validate the Visual Recognition Assessment–Leuven (ViRA-L), a recognition-memory battery assessing ten visual categories within a matched task structure, and provide age-stratified normative data.

**Method:** Healthy Dutch-speaking adults completed a visual recognition task for ten categories. Each category included six targets, 18 Learning trials, and 30 Novel trials requiring recognition across appearance changes. After exclusions, 417 participants remained. We evaluated reliability, score distributions, task effects, demographic associations, and category-specific norms.

**Results:** Battery-level internal consistency was good (Cronbach’s α =.879; omega total =.901), with lower category-level estimates, broad score dispersion, and minimal floor and ceiling effects. Novel and difficult trials produced lower accuracy and slower responses. Age was the clearest demographic correlate of accuracy; gender and education showed no independent overall associations. Age effects varied by category and were larger in Novel than Learning trials. Model-based differences between ages 50 and 85 were largest for Pseudowords and Hands and smallest for Houses.

**Conclusions:** The ViRA-L supports comparison of recognition-memory profiles under matched procedures. Category-dependent age associations support age-sensitive, category-specific interpretation. The battery provides a foundation for neuropsychological research; further validation is required for clinical diagnostic use.

**Statement of Research Significance:** *Research Question(s) or Topic(s):* Can one recognition-memory task compare performance across ten visual categories, and how do these abilities differ between younger and older adults?

*Main Findings:* The battery produced reliable overall scores, while reliability varied between categories. Older age was associated with lower accuracy, particularly for pseudowords and hands, and when learned items appeared in new views. Age differences were smaller for houses.

*Study Contributions:* The study provides a common framework and age-specific reference scores for examining recognition profiles across categories. It supports future research on visual difficulties after brain injury and identifies the additional validation needed before diagnostic use.

## Introduction

Visual recognition enables the identification of faces, words, and objects despite changes in viewpoint, illumination, and image quality. Damage to ventral occipitotemporal cortex can disrupt this ability, producing prosopagnosia, pure alexia, or object agnosia (Bornstein & Kidron, 1959; Collignon, 1972; James et al., 2003; Konen et al., 2011). These disorders have informed debates about specialization in high-level vision, yet patients commonly show deficits spanning several domains (Behrmann & Plaut, 2020; Geskin & Behrmann, 2018; Rice et al., 2021). Mixed profiles may reflect partly shared processing mechanisms or damage to several specialized systems and their connections. An assessment sampling multiple categories can characterize these profiles without presupposing either architecture.

Neuroimaging similarly reveals specialization within a distributed system. Regions respond preferentially to faces, bodies, written words, and scenes (Cohen et al., 2000; Epstein & Kanwisher, 1998; Kanwisher et al., 1997; Taylor et al., 2007), while overlapping activity patterns carry information about multiple categories (Bracci & Op de Beeck, 2016; Haxby et al., 2001; Kriegeskorte et al., 2008). Occipitotemporal organization also follows broader dimensions, including animacy, real-world size, eccentricity, and behaviorally relevant object properties (Contier et al., 2024; Hasson et al., 2002; Konkle & Oliva, 2012; Op de Beeck et al., 2019; Sha et al., 2015). Understanding recognition therefore requires determining how specialized and shared mechanisms jointly shape performance across domains.

Behavioral comparisons are complicated by differences between assessment tasks. Faces are commonly assessed through unfamiliar-face memory, words through lexical decisions or reading, and objects through naming or matching. Apparent associations or dissociations may consequently reflect memory load, response format, decision demands, or stimulus transformations rather than category-specific visual abilities (Behrmann & Plaut, 2020; Robotham & Starrfelt, 2017). The Cambridge Car Memory Test addressed this problem by matching the format of the Cambridge Face Memory Test (CFMT; Dennett et al., 2012; Duchaine & Nakayama, 2006). The Vanderbilt Expertise Test extended a common framework across several object categories (McGugin et al., 2012). These advances motivate a battery that combines matched procedures with broader coverage of categories relevant to neuropsychology and cortical organization.

Matched procedures matter in neuropsychology, where selective impairment is inferred from contrasts between affected and relatively preserved abilities. A demanding face-memory test and an easier object-naming task could suggest a dissociation partly because they differ in memory and decision requirements. Conversely, shared task demands may contribute to correlated scores. Sampling several categories within one procedure allows these possibilities to be examined and reduces dependence on a single comparison category. This does not eliminate differences between domains, but makes their procedural basis explicit (Behrmann & Plaut, 2020; Robotham & Starrfelt, 2017).

We developed the Visual Recognition Assessment–Leuven (ViRA-L). Its ten categories are Cars, Bodies, Musical Instruments, Pseudowords, Tools, Faces, Food, Houses, Hands, and Animals. Faces and pseudowords address clinically prominent abilities with well-characterized cortical selectivity and a debated functional relationship (Cohen et al., 2000; Dehaene et al., 2010; Robotham & Starrfelt, 2017). Bodies and Hands sample distinctions within body-related processing (Bracci et al., 2010; Downing et al., 2001; Spiridon et al., 2006). Houses represent the scene domain, while Tools and Food extend coverage to manipulable objects and biologically relevant consumables (Epstein & Kanwisher, 1998; Khosla et al., 2022; Martin et al., 1996). Collectively, the categories also sample animacy and real-world size, major dimensions of occipitotemporal organization (Ritchie et al., 2021; Sha et al., 2015). Thus, selection was guided by lesion relevance and cortical organization rather than by the assumption that each category corresponds to an independent module. Broader coverage allows relationships among several affected and spared abilities to be examined within the same individual.

Matched assessment additionally requires appropriate normative references. A category score can only be interpreted relative to the distribution expected in comparable healthy individuals (Karstens et al., 2024; Lezak, 2004; Mitrushina, 2005). This is especially relevant across adulthood, when perceptual acuity, memory, and processing speed change at different rates (Hedden & Gabrieli, 2004; van Hooren et al., 2007). Aging can delay high-level visual representations and reduce their selectivity, with potentially different consequences across category systems (Bourbon-Teles et al., 2021; Haupt et al., 2025; Koen & Rugg, 2019). A common age-aggregated benchmark could therefore obscure atypical performance in one domain while overinterpreting age-typical performance in another. Category-specific, age-sensitive norms are needed alongside comparable task demands (Devora et al., 2020; Wang et al., 2021; Wragg et al., 2024).

The ViRA-L applies Cambridge-style recognition-memory procedures across all ten categories. Participants learn target identities and identify them among distractors under changed viewing conditions; performance therefore involves perceptual discrimination and memory. This study establishes battery-and category-level reliability and score dispersion, evaluates the expected effects of task stage and difficulty, examines demographic and category-specific age associations, and provides age-stratified normative data. These analyses establish a basis for comparing recognition profiles across domains and for subsequent investigation of neurological impairment.

## Method

### Participants and procedure

We recruited 439 native Dutch-speaking adults, comprising 339 students and 100 older adults, who reported normal or corrected-to-normal vision and no neurological disease. Students were recruited through Sona Systems; older adults through a volunteer database, personal contacts, and social media. All provided written informed consent. The study followed the Declaration of Helsinki and received local Ethics Committee approval by the Social and Societal Ethics Committee (G-2023-6837-R2(AMD)). Students received course credit and older adults €12 per hour.

After preprocessing, 417 participants remained: 322 students and 95 older adults. Of 61 students administered randomized category order, 57 remained and contributed only to order-effect analyses. The primary fixed-order sample comprised 360 participants, including 265 students and 95 older adults. Table 1 describes the final sample.

**Table 1.** Demographic characteristics of the final sample.

| **Variable** | **Total** | **Students** | **Older adults** |
|---|---|---|---|
| N | 417 | 322 | 95 |
| Mean age in years (SD, min–max) | 29.58 (21.18, 18–85) | 18.42 (1.39, 18–35) | 68.16 (8.31, 50–85) |
| Education, % low | 6.04 | 0 | 27.17 |
| Education, % high | 93.96 | 100 | 72.83 |
| Gender, % male | 20.87 | 15.62 | 39.13 |
| Handedness, % right | 87.8 | 88.2 | 86.3 |
| Handedness, % left | 10.3 | 11.8 | 5.3 |
| Handedness, % ambidextrous | 1.2 | 0.0 | 5.3 |
| MoCA completed, *n* (of which Blind MoCA) | 50 (29) | 0 (0) | 50 (29) |

Note. Demographic characteristics are shown for the final sample after preprocessing and participant-level exclusions. Education level was dichotomized into low (completed secondary education or less) and high (any education pursued beyond secondary education). Missing data were observed for age (*N* = 7; of which 3 older adults), education (*N* = 3 older adults), gender (*N* = 5, of which 3 older adults), and handedness (*N* = 3 older adults). MoCA counts refer to the final older-adult sample; a score out of 22 denotes the Blind MoCA used for telephone administration.

Participants completed a demographic questionnaire and then the battery. Nine completed it on site with an experimenter; the remainder participated remotely. Remote participants were instructed to use a computer in a quiet environment, maintain a comfortable viewing distance, and avoid interruptions. Everyone completed the battery within one day. Students completed one continuous task; older adults completed two parts with an optional break. Screen characteristics, viewing distance, distractions, compliance, and break duration were not objectively monitored.

Fifty-five older adults completed the Montreal Cognitive Assessment (MoCA; Nasreddine et al., 2005) after the ViRA-L: 32 received the telephone-administered 22-point version and 23 the 30-point version on site. For telephone administration, visually presented MoCA items were omitted. Screening required a separate appointment, which some participants did not arrange or attend. After exclusions, 50 completers remained, including 29 telephone and 21 full-version administrations.

### Task and stimuli

The task was programmed in PsychoPy Builder and deployed through Pavlovia (Open Science Tools Ltd., 2026; Peirce et al., 2019). Each category used the same target count, trial structure, and three-alternative forced-choice response format (Figure 1). Study images were unavailable during recognition trials, requiring memory for previously studied identities rather than simultaneous perceptual matching.

**Figure 1.**
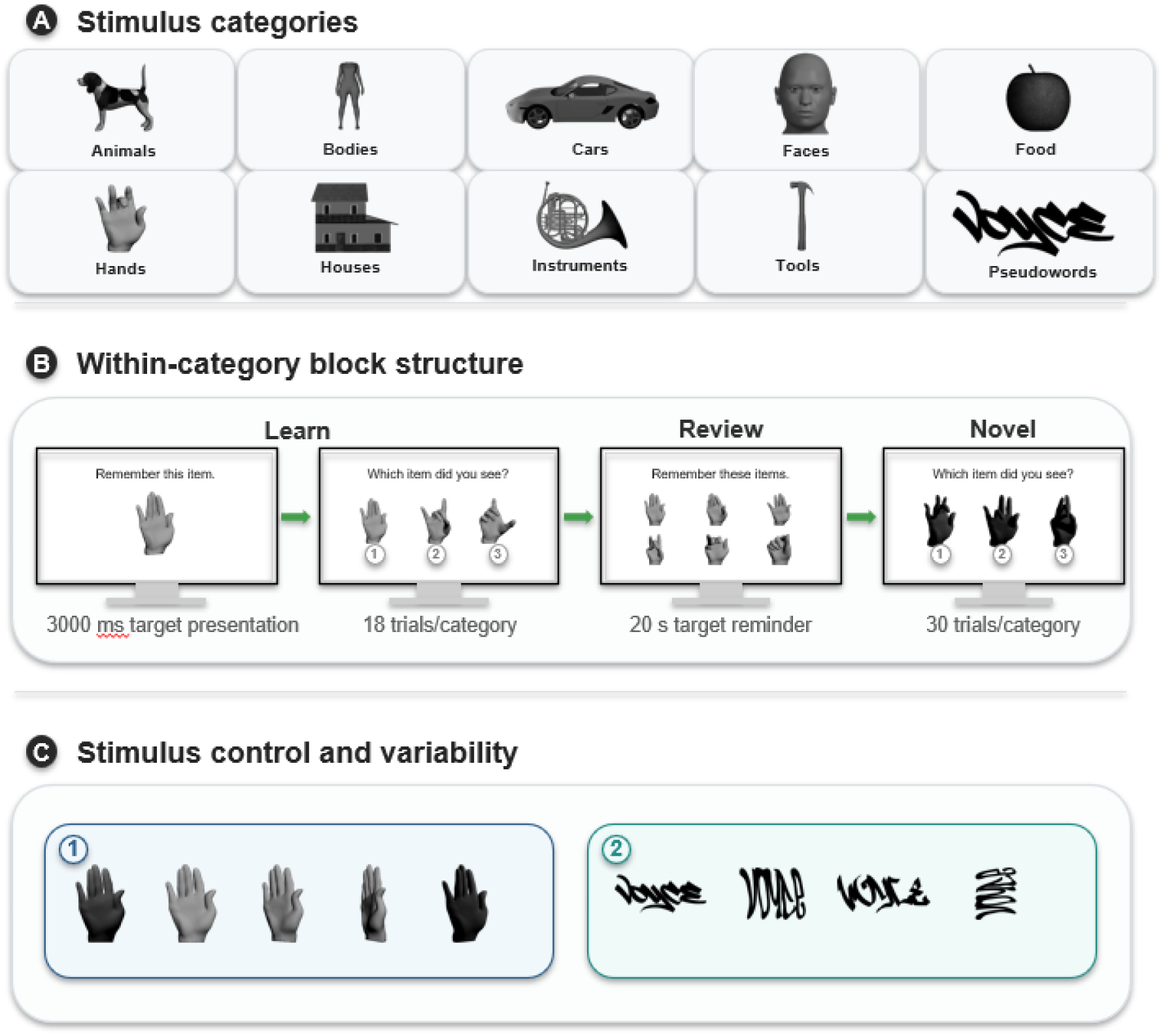
Overview of the ViRA-L (Visual Recognition Assessment – Leuven) Note. A. The battery comprises ten visual categories, each tested in a separate block, with representative exemplars shown for each category. B. Each block followed an identical recognition-memory sequence modelled on the Cambridge Face Memory Test. In the Learning Stage, a target was presented for 3000 ms and followed by three-alternative forced-choice (3AFC) recognition trials (18 per category); a review screen then re-presented the targets for 20 s, after which the Novel Stage probed recognition among new distractors (30 trials per category). Trials were separated by a 500 ms interstimulus interval. C. To ensure recognition required fine-grained discrimination rather than low-level image cues, each target was shown across multiple viewpoints and lighting conditions (panel 1); for Pseudowords, visual form was instead varied by changing the font (panel 2). Stimulus sources for all ten categories are detailed in Supplementary Table S1. For illustrative purposes, photographic face stimuli were replaced with computer-generated exemplars in this figure; the original photographs were used during experimental testing.

Each block began with one practice target presented in three views and three practice trials with feedback. During the Learning Stage, each of six target identities appeared in three study images, each shown for 3,000 ms with 500-ms intervals, followed by three recognition trials. Repeating this sequence yielded 18 Learning trials. All six targets were then shown in a 2 × 3 review display for 20 s. The Novel Stage comprised 30 trials, each presenting one learned target and two distractors in one of six newly generated views differing from studied images in orientation, illumination, or both. Each category contributed 48 scored trials. Participants responded with number keys corresponding to the top, middle, and bottom images, using their preferred hand. Displays remained until response; intertrial intervals were 500 ms.

Older adults received five additional motivational easy trials per category during the Novel Stage. These paired a learned target with visually dissimilar distractors, all in frontal view, and were excluded from scoring. Standard administration fixed category and within-category order; the order-control version randomized category blocks while preserving within-block order. The complete battery required approximately 60–80 minutes. We omitted the CFMT noise stage to limit duration, supported by evidence that the first 48 CFMT items retain performance characteristics comparable to the full test (Corrow et al., 2018).

Each category contained six targets and 30 unique distractors, yielding 96 distractor presentations. Controlled distractor repetition reduced the usefulness of simple familiarity judgments. Relatively homogeneous subcategories increased within-category similarity and discouraged coarse semantic labeling. Transformations were adapted to object geometry because identical rotation axes would not produce comparable perceptual changes across categories. Image-based easy and difficult trials used lower-and higher-similarity target–distractor pairs, respectively, quantified with the Structural Similarity Index Measure (SSIM; Wang et al., 2004). Pairwise grayscale similarity scores were ranked within category and divided into upper and lower halves to construct difficult and easy pairings. Pseudoword difficulty instead depended on orthographic similarity. Item assignment and stimulus parameters were refined during piloting to obtain substantial discrimination demands without pronounced floor or ceiling effects. Stimulus sources, subcategories, transformations, and trial construction appear in Supplementary Materials S1-S2.

Pseudowords preserved legal orthographic structure while removing lexical meaning, reducing reliance on vocabulary and stored word knowledge and facilitating future language adaptation. They were generated with Wuggy (Keuleers & Brysbaert, 2010); fonts varied appearance in place of lighting transformations. All stimuli were converted to grayscale, resized while preserving aspect ratio, and adjusted to reduce extreme brightness differences without removing internal luminance structure. Images were displayed centrally against dark gray in normalized PsychoPy units. Their size was standardized relative to the browser window, but not in physical dimensions or visual angle. Source dimensions and additional administration details are provided in Supplementary Material S4.1.

### Preprocessing and scoring

Analyses used R 4.5.2 (R Core Team, 2025). Prespecified trial exclusions removed one observation with missing accuracy or reaction time, 5,140 additional easy trials, and 1,361 duplicate rows. Duplicates arose when remote participants repeated task sections because they were unsure whether responses had been saved; the first occurrence was retained. Responses exceeding each participant’s mean reaction time by three standard deviations removed 3,335 trials. Lower thresholds were estimated separately by cohort from the earliest stable crossover at which the log-reaction-time density for correct responses exceeded that for errors (Crouzet et al., 2014; Voormann & Miller, 2024; Voskuilen et al., 2018). Thresholds of 591 ms for students and 739 ms for older adults removed 6,405 trials, leaving 200,914. Alternative smoothing yielded similar thresholds (Supplementary Material S3.1).

Sixteen participants were excluded for retaining fewer than 60% of trials in at least four categories, and two for below-chance performance in more than one category. Four older adults scored below the ninth percentile on version-appropriate MoCA norms: Dutch age-, sex-, and education-adjusted norms for the full version (Kessels et al., 2022) and demographically adjusted MoCA-22 norms (Ratcliffe et al., 2024). The final dataset comprised 194,635 trials. Category accuracy was the percentage correct across valid trials; total accuracy used all valid trials. Reaction-time analyses included correct responses only, log-transformed before modeling. Additional exclusion counts by cohort, duplicate-identification rules, and scoring definitions appear in Supplementary Materials S4.2–S4.3.

### Statistical analyses and normative data

Psychometric analyses used the fixed-order sample. Cronbach’s *α*, omega total, and Spearman–Brown-corrected odd–even split-half reliability were calculated across ten category scores for the battery and six items within each category. Item difficulty was the proportion correct for each item across participants; higher values therefore indicated easier items. Corrected item–total correlations related each item to the mean of the other items in its category, excluding the item itself from the comparison score. These indices assessed item functioning separately from overall score dispersion. Distributional summaries included mean, standard deviation, range, skewness, and Shapiro–Wilk tests. Floor and ceiling effects were defined as scores at or below chance (33.3%) and at 100%, respectively.

Trial-level accuracy was analyzed using binomial generalized linear mixed-effects models with participant and item random intercepts. Fixed effects were age, gender, education, stage, category, and difficulty. Corresponding linear mixed-effects models analyzed correct-trial log reaction time. Accuracy coefficients used Wald z tests; reaction-time coefficients used t tests with Satterthwaite degrees of freedom. Estimated marginal means compared Learning and Novel stages within categories with Holm correction. Participant-level Pearson correlations related total accuracy to mean reaction time and its within-participant standard deviation. Logistic mixed models compared fixed and randomized category order.

Age was modeled linearly or using natural cubic splines with three or four degrees of freedom; Akaike information criterion selected the three-degree specification. Likelihood-ratio tests assessed age’s overall contribution and interactions with category, difficulty, and stage. Category-specific models followed the Age × Category interaction. Estimated marginal means summarized nonlinear age associations as contrasts between ages 50 and 85. Orientation was assessed separately within stages because available transformations differed; models with and without orientation were compared by likelihood-ratio tests. Within-category subcategory effects were tested with false-discovery-rate correction across ten tests. Supplementary Material S4.4 provides further model specifications.

Norms used first-session fixed-order data and category scores collapsed across Learning and Novel trials; no total-battery norms were generated. Four participants without age information were excluded, leaving 356. For ages 18–49, 50–64, 65–74, and 75+, we report sample size, mean, standard deviation, and the 5th, 16th, 25th, 50th, 75th, and 95th percentiles. Cells below 30 participants were flagged. Age was the clearest demographic correlate; further stratification would fragment the sample. Grouped empirical norms avoided continuous regression-based interpolation across the unsampled interval between younger and older cohorts (Supplementary Material S4.5).

## Results

### Psychometric properties

Battery-level internal consistency was good in the fixed-order sample (*N* = 360): α =.879, 95% CI [.860,.897], omega total =.901, and split-half reliability =.887. Category estimates were lower, with *α* ranging from.466 to.756, omega from.674 to.823, and split-half reliability from.480 to.793; Cars and Pseudowords defined the lower and upper limits. Mean corrected item–total correlations similarly ranged from.253 for Cars to.525 for Pseudowords (Supplementary Tables S1–S2).

Total accuracy was broadly distributed (*M* = 78.69%, *SD* = 7.01, range = 54.58–95.91), with slight negative skew (−0.48). Category means ranged from 65.22% for Cars to 86.68% for Animals; Houses also scored relatively highly (84.69%), whereas Faces scored 71.99%. Means for the remaining categories were Food, 80.65%; Musical Instruments, 80.07%; Bodies, 79.91%; Pseudowords, 79.50%; Hands, 79.03%; and Tools, 78.57%. Floor and ceiling scores were uncommon, and no participant achieved a perfect total score. Supplementary Table S3 reports all category distributions; exact floor and ceiling counts are retained in Supplementary Material S5.1. Fixed and randomized category orders did not differ in overall accuracy (*OR* = 1.036, 95% CI [0.902, 1.190], *p* =.619), and category profiles were similar across versions (Figure 2). Thus, the observed category pattern was not accompanied by a detectable overall administration-order effect, although this comparison does not establish equivalence between versions.

**Figure 2.**
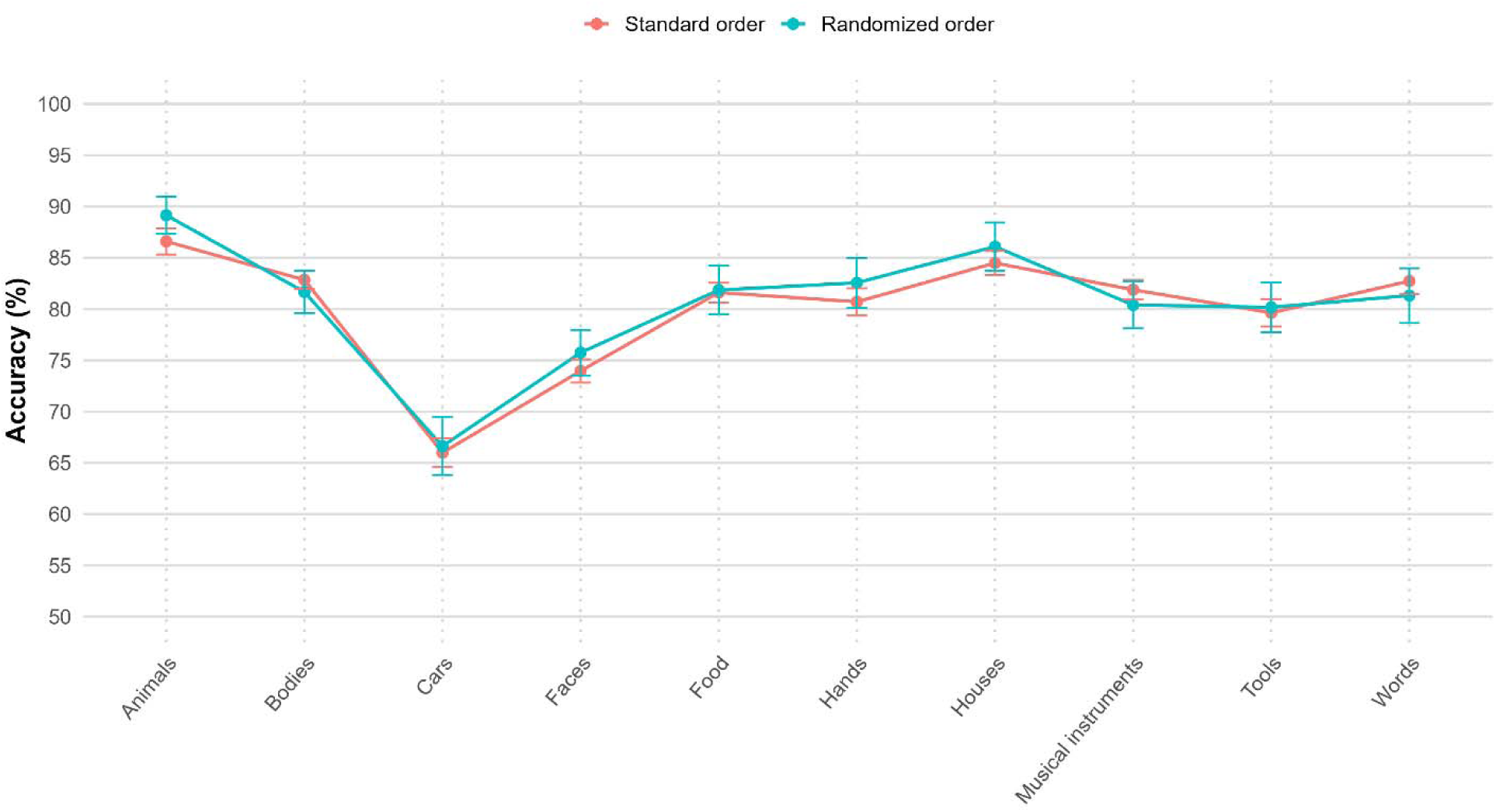
Accuracy by category in the standard and randomized versions of the task. Note. Points show mean accuracy and error bars denote 95% confidence intervals for the standard fixed-order and randomized-order versions.

### Task characteristics

Accuracy was lower for Novel (*M* = 70.64%, *SD* = 9.19) than Learning trials (*M* = 91.87%, *SD* = 4.28; *OR* = 0.169, 95% CI [0.163, 0.175], *p* <.001). Stage effects varied by category, χ²(9) = 818.26, *p* <.001, but every category showed lower Novel accuracy after Holm correction (Figure 3; Supplementary Table S9). Difficult trials likewise produced lower accuracy (*M* = 75.08%, *SD* = 8.06) than easy trials (*M* = 82.34%, *SD* = 6.33; *OR* = 0.571, 95% CI [0.555, 0.587], *p* <.001).

**Figure 3.**
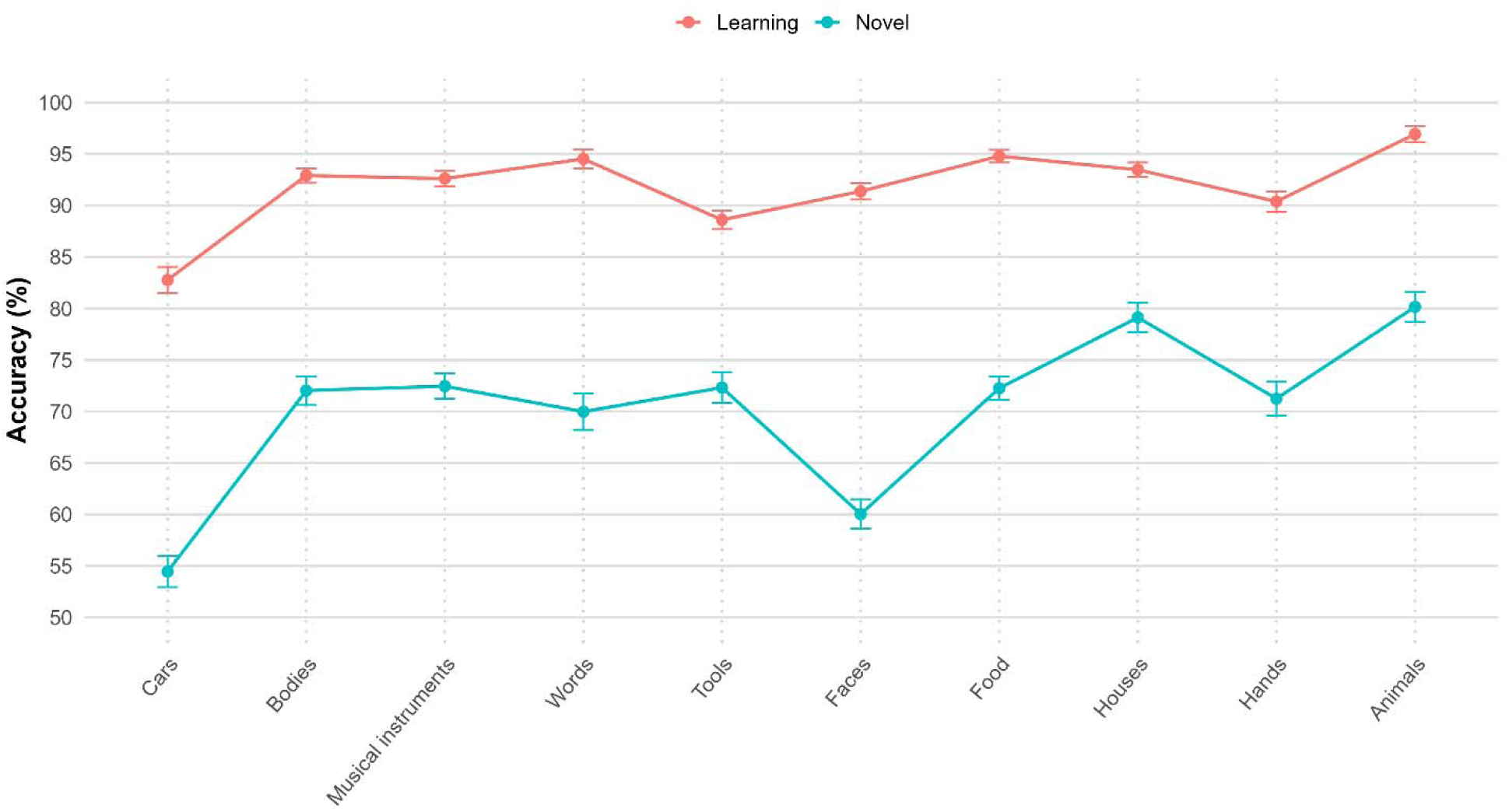
Performance by category in the Learning and Novel Stages. Note. Points show mean category-level accuracy in the fixed-order primary sample; error bars indicate 95% confidence intervals computed across participants.

Responses were slower for Novel (*M* = 3.94 s, *SD* = 1.70) than Learning trials (*M* = 2.64 s, *SD* = 1.05; β = 0.463, 95% CI [0.457, 0.469], *p* <.001), and for difficult (*M* = 3.62 s, *SD* = 1.52) than easy trials (*M* = 3.13 s, *SD* = 1.25; *β* = 0.161, 95% CI [0.156, 0.166], *p* <.001). These coefficients correspond to geometric-mean reaction-time ratios of 1.589 and 1.175. Accuracy showed little association with mean reaction time, *r*(358) = −.042, 95% CI [−.145,.062], *p* =.428, or its variability, *r*(358) =.054, 95% CI [−.050,.156], *p* =.308.

Orientation improved model fit in Learning, χ²(2) = 98.99, and Novel trials, χ²(4) = 346.86, both *p* <.001. Learning estimates remained high across frontal and 30° views (95.78%–96.96%). Novel estimates ranged from 81.58% for frontal views with altered shadows to 74.79% for rotation with lateral shadow; 60° rotations without shadow were intermediate (supplementary Tables S10A and S14). Viewpoint and illumination were not fully crossed, so their independent contributions cannot be isolated. Subcategory effects were nominally significant for Bodies, Pseudowords, and Food, but none survived false-discovery-rate correction (Supplementary Table S6).

### Demographic associations

Age predicted accuracy, χ²(3) = 49.39, *p* <.001. The overall age-85-minus-age-50 contrast was −14.5 percentage points (95% CI [−19.9, −9.0], *OR* = 0.370, 95% CI [0.260, 0.526], both *p* <.001). Overall associations were nonsignificant for gender (male versus female: *OR* = 1.008, 95% CI [0.890, 1.141], *p* =.906) and education (higher versus lower: *OR* = 1.160, 95% CI [0.950, 1.418], *p* =.145). The age result was similar before MoCA exclusions, χ²(3) = 50.29, *N* = 358, *p* <.001, and after exclusions, *N* = 355 (Supplementary Material S3.4).

Age associations differed across categories (χ²(27) = 344.47, *p* <.001 (Figure 4)). Predicted age-85-minus-age-50 accuracy differences were largest for Pseudowords (−23.26 percentage points), followed by Hands (−20.45), Cars (−17.84), Bodies (−17.78), Musical Instruments (−17.60), Faces (−14.52), Tools (−9.99), Food (−9.27), Animals (−8.45), and Houses (−6.68). Percentage-point CIs were [−31.8, −14.7] for Pseudowords, [−28.6, −12.3] for Hands, and [−11.4, −1.9] for Houses; all category contrasts were significant (Supplementary Table S4).

**Figure 4.**
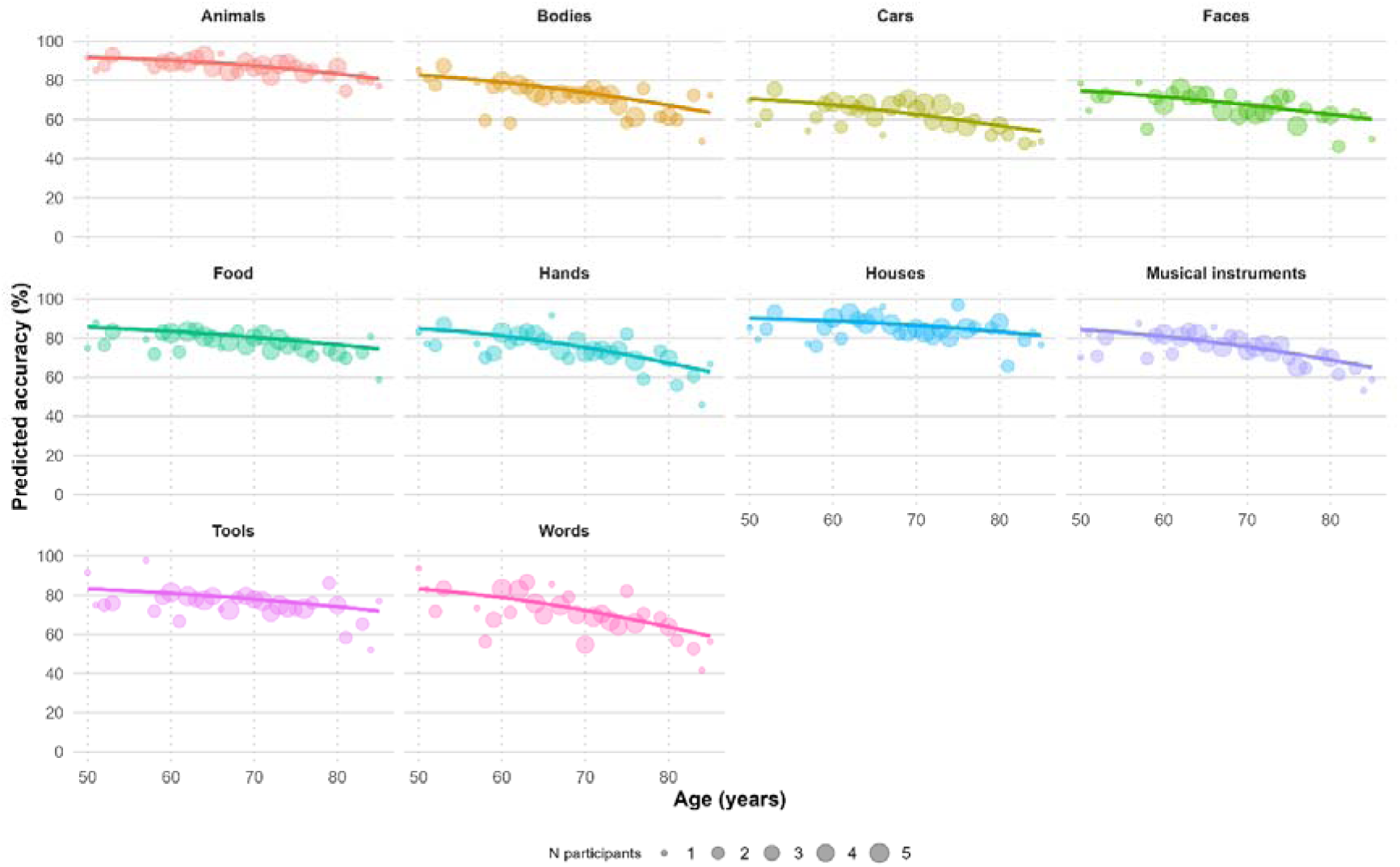
Category-specific age effects. Note. Lines show model-based predicted accuracy by age for each category, adjusted for gender and education and averaged over stage and difficulty. Points show observed mean accuracy at each age, with bubble size indicating the number of participants contributing to that value.

Age interacted with stage, χ²(3) = 8.78, *p* =.032, but not difficulty, χ²(3) = 1.17, *p* =.760. Predicted differences were −21.79 percentage points for Novel, 95% CI [−29.6, −13.93], and −7.89 for Learning, 95% CI [−11.1, −4.63]. Their difference was −13.9 points, 95% CI [−19.4, −8.36], *p* <.001; the corresponding ratio of odds ratios was nonsignificant (1.11, 95% CI [0.886, 1.39], *p* =.365), indicating scale-dependent interaction estimates. Reaction time varied with age, χ²(3) = 285.79, and showed Age × Stage, χ²(3) = 345.45, and Age × Difficulty interactions, χ²(3) = 28.38, all *p* <.001. The age-85-versus-age-50 reaction-time ratio was 1.43, 95% CI [1.19, 1.71], *p* <.001. Higher education was associated with slower responses (*β* = 0.142, *SE* = 0.052, 95% CI [0.040, 0.243], *p* =.007); gender was not (*β* = −0.026, 95% CI [−0.088, 0.036], *p* =.413). Complete coefficients and follow-ups appear in Supplementary Tables S5 and S7–S10.

### Normative data

Age-band sample sizes were 264 for ages 18–49, 33 for 50–64, 38 for 65–74, and 21 for 75+ (Table 2). The first three met the cell-size criterion; the oldest requires caution. The nominal 18–49 band contained participants aged 18–35 only. Their total accuracy (*M* = 80.09%, *SD* = 6.60) did not differ significantly from the 50–64 group (*M* = 78.70%, *SD* = 5.49; difference = 1.39 percentage points, 95% CI [−0.70, 3.48], *t*(44.44) = 1.34, *p* =.188; Hedges’ g = 0.21, 95% CI [−0.15, 0.57]). This comparison does not establish performance at the unobserved ages 36–49.

**Table 2.**
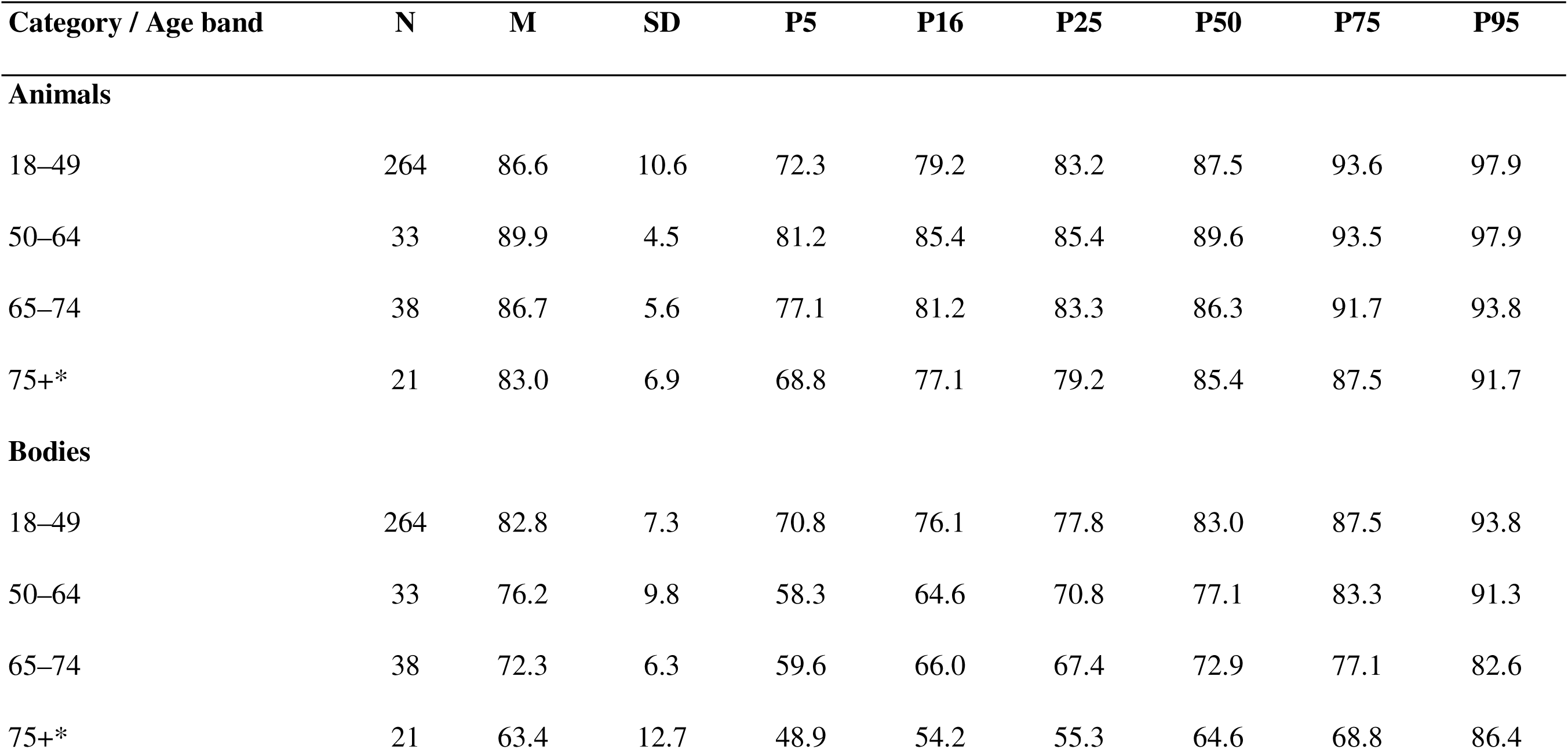
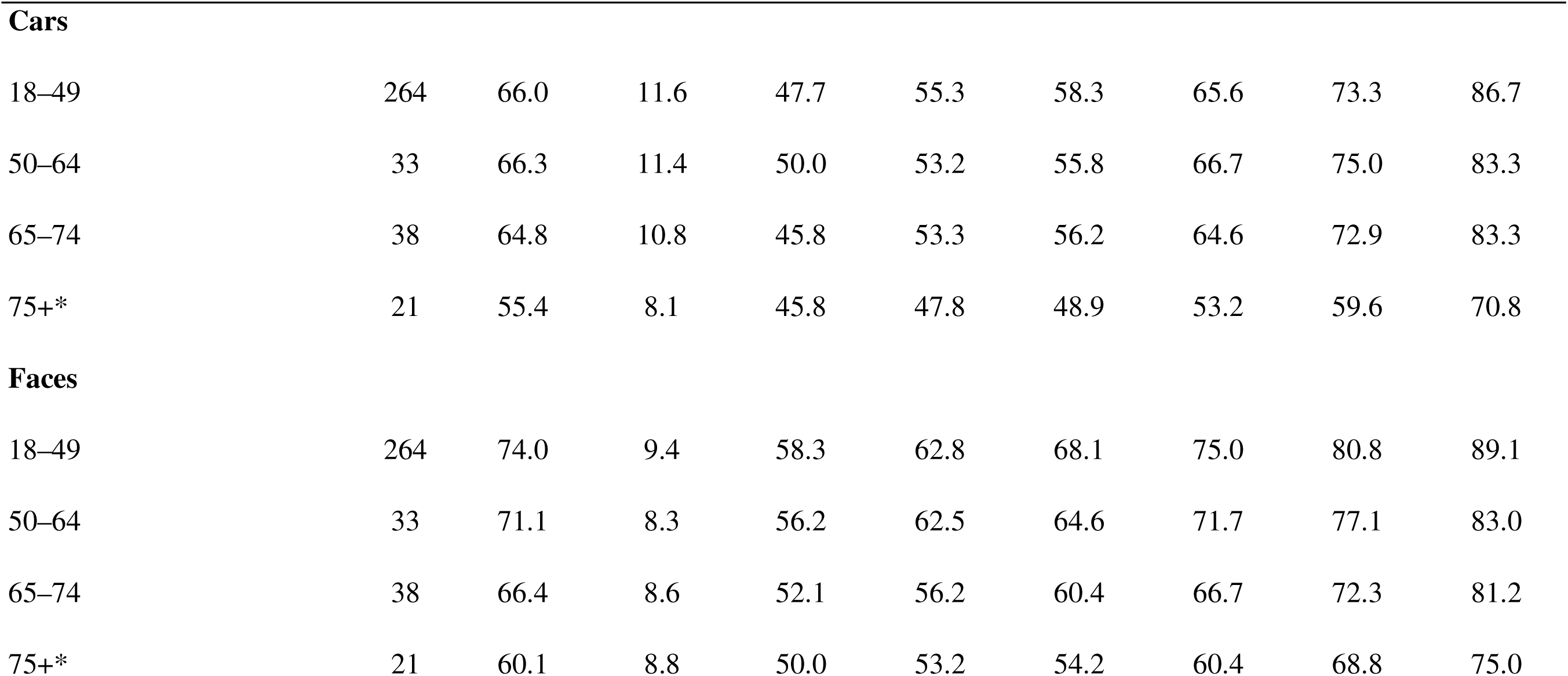
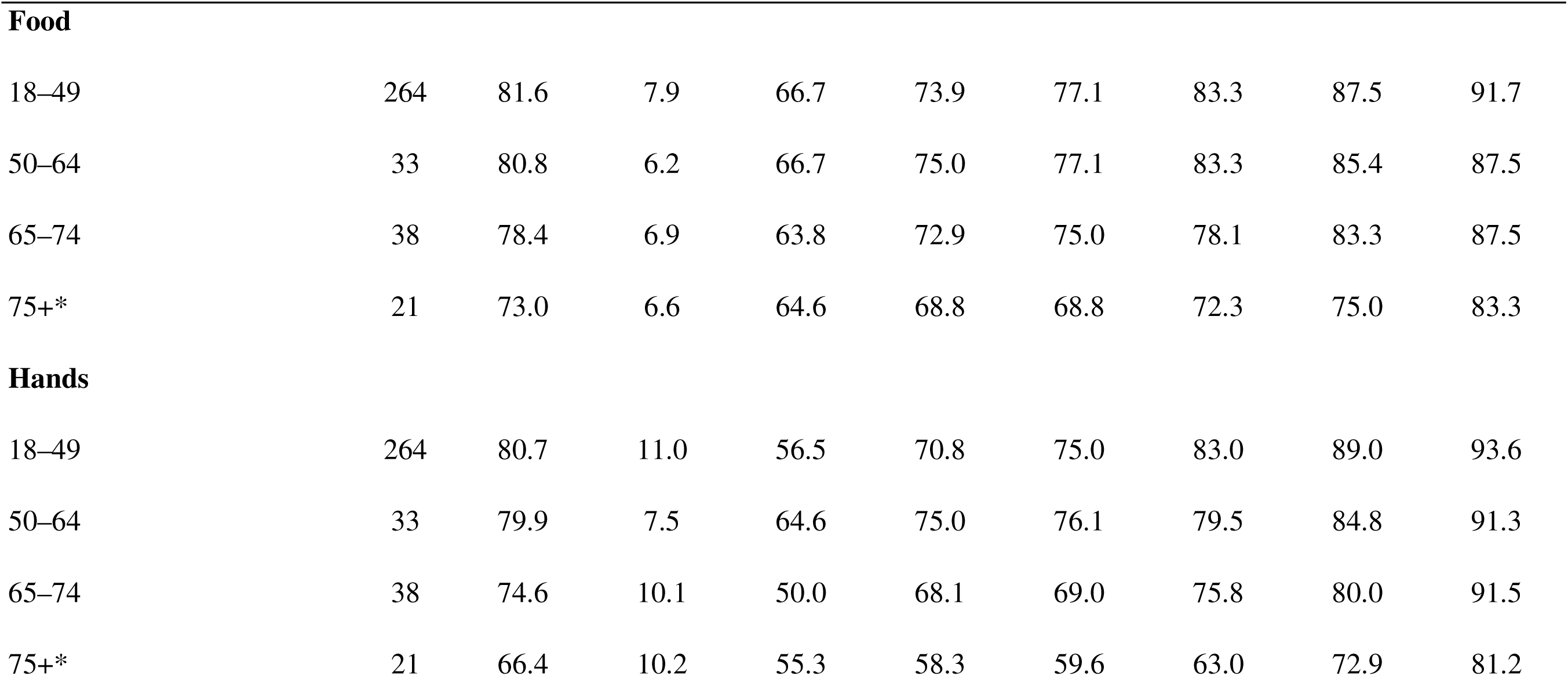
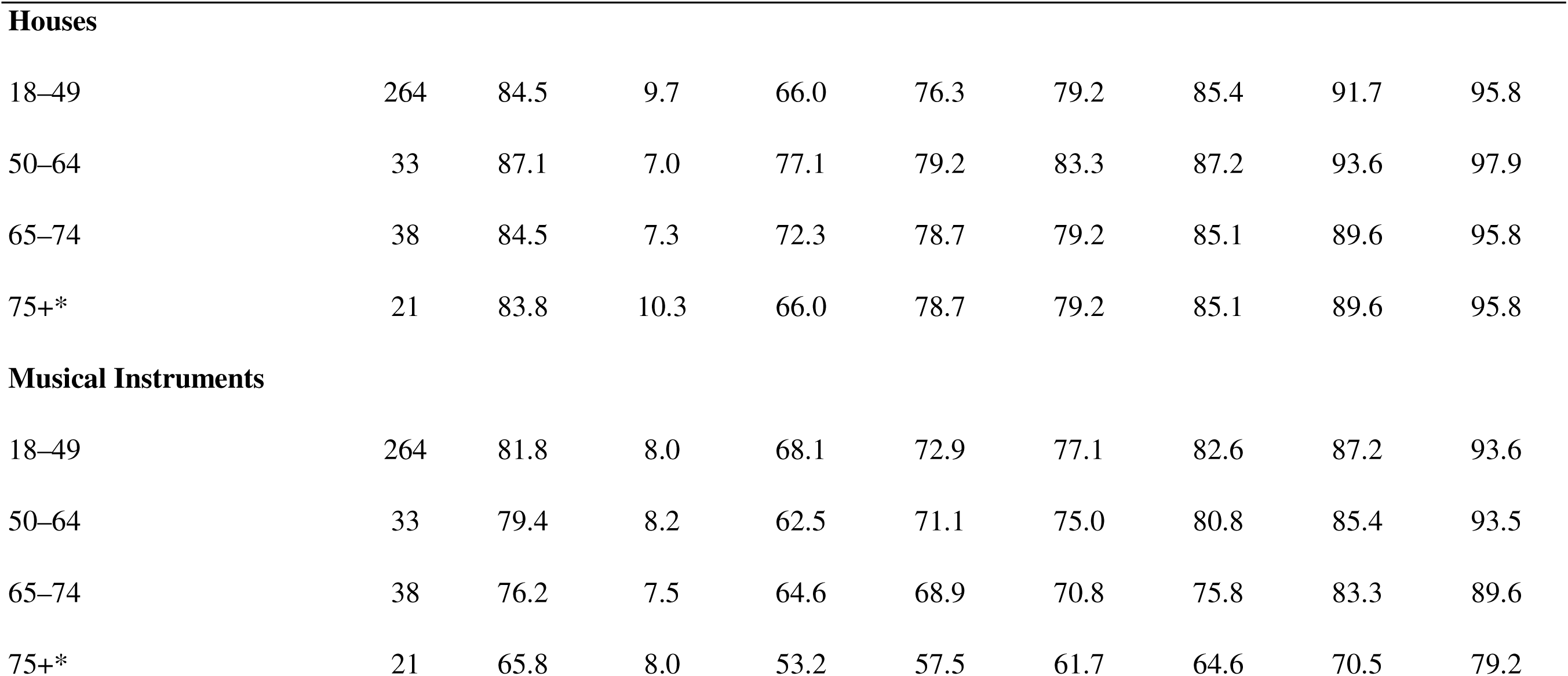
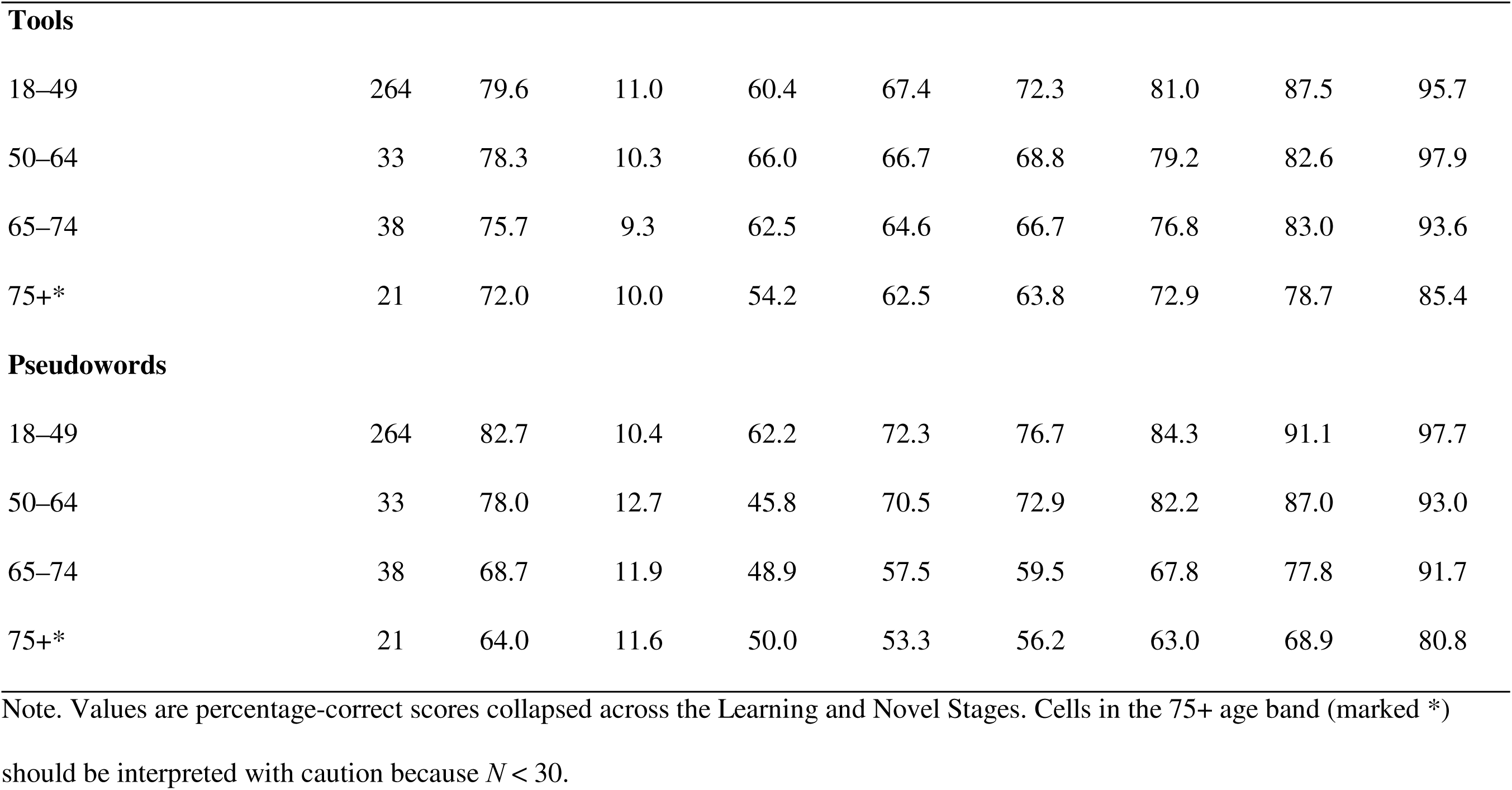
Category-level normative scores by age band for the ViRA-L (%)

| **Category / Age band** | **N** | **M** | **SD** | **P5** | **P16** | **P25** | **P50** | **P75** | **P95** |
|---|---|---|---|---|---|---|---|---|---|
| **Animals** |  |  |  |  |  |  |  |  |  |
| 18–49 | 264 | 86.6 | 10.6 | 72.3 | 79.2 | 83.2 | 87.5 | 93.6 | 97.9 |
| 50–64 | 33 | 89.9 | 4.5 | 81.2 | 85.4 | 85.4 | 89.6 | 93.5 | 97.9 |
| 65–74 | 38 | 86.7 | 5.6 | 77.1 | 81.2 | 83.3 | 86.3 | 91.7 | 93.8 |
| 75+* | 21 | 83.0 | 6.9 | 68.8 | 77.1 | 79.2 | 85.4 | 87.5 | 91.7 |
| **Bodies** |  |  |  |  |  |  |  |  |  |
| 18–49 | 264 | 82.8 | 7.3 | 70.8 | 76.1 | 77.8 | 83.0 | 87.5 | 93.8 |
| 50–64 | 33 | 76.2 | 9.8 | 58.3 | 64.6 | 70.8 | 77.1 | 83.3 | 91.3 |
| 65–74 | 38 | 72.3 | 6.3 | 59.6 | 66.0 | 67.4 | 72.9 | 77.1 | 82.6 |
| 75+* | 21 | 63.4 | 12.7 | 48.9 | 54.2 | 55.3 | 64.6 | 68.8 | 86.4 |
| **Cars** |  |  |  |  |  |  |  |  |  |
| 18–49 | 264 | 66.0 | 11.6 | 47.7 | 55.3 | 58.3 | 65.6 | 73.3 | 86.7 |
| 50–64 | 33 | 66.3 | 11.4 | 50.0 | 53.2 | 55.8 | 66.7 | 75.0 | 83.3 |
| 65–74 | 38 | 64.8 | 10.8 | 45.8 | 53.3 | 56.2 | 64.6 | 72.9 | 83.3 |
| 75+* | 21 | 55.4 | 8.1 | 45.8 | 47.8 | 48.9 | 53.2 | 59.6 | 70.8 |
| **Faces** |  |  |  |  |  |  |  |  |  |
| 18–49 | 264 | 74.0 | 9.4 | 58.3 | 62.8 | 68.1 | 75.0 | 80.8 | 89.1 |
| 50–64 | 33 | 71.1 | 8.3 | 56.2 | 62.5 | 64.6 | 71.7 | 77.1 | 83.0 |
| 65–74 | 38 | 66.4 | 8.6 | 52.1 | 56.2 | 60.4 | 66.7 | 72.3 | 81.2 |
| 75+* | 21 | 60.1 | 8.8 | 50.0 | 53.2 | 54.2 | 60.4 | 68.8 | 75.0 |
| **Food** |  |  |  |  |  |  |  |  |  |
| 18–49 | 264 | 81.6 | 7.9 | 66.7 | 73.9 | 77.1 | 83.3 | 87.5 | 91.7 |
| 50–64 | 33 | 80.8 | 6.2 | 66.7 | 75.0 | 77.1 | 83.3 | 85.4 | 87.5 |
| 65–74 | 38 | 78.4 | 6.9 | 63.8 | 72.9 | 75.0 | 78.1 | 83.3 | 87.5 |
| 75+* | 21 | 73.0 | 6.6 | 64.6 | 68.8 | 68.8 | 72.3 | 75.0 | 83.3 |
| **Hands** |  |  |  |  |  |  |  |  |  |
| 18–49 | 264 | 80.7 | 11.0 | 56.5 | 70.8 | 75.0 | 83.0 | 89.0 | 93.6 |
| 50–64 | 33 | 79.9 | 7.5 | 64.6 | 75.0 | 76.1 | 79.5 | 84.8 | 91.3 |
| 65–74 | 38 | 74.6 | 10.1 | 50.0 | 68.1 | 69.0 | 75.8 | 80.0 | 91.5 |
| 75+* | 21 | 66.4 | 10.2 | 55.3 | 58.3 | 59.6 | 63.0 | 72.9 | 81.2 |
| **Houses** |  |  |  |  |  |  |  |  |  |
| 18–49 | 264 | 84.5 | 9.7 | 66.0 | 76.3 | 79.2 | 85.4 | 91.7 | 95.8 |
| 50–64 | 33 | 87.1 | 7.0 | 77.1 | 79.2 | 83.3 | 87.2 | 93.6 | 97.9 |
| 65–74 | 38 | 84.5 | 7.3 | 72.3 | 78.7 | 79.2 | 85.1 | 89.6 | 95.8 |
| 75+* | 21 | 83.8 | 10.3 | 66.0 | 78.7 | 79.2 | 85.1 | 89.6 | 95.8 |
| **Musical Instruments** |  |  |  |  |  |  |  |  |  |
| 18–49 | 264 | 81.8 | 8.0 | 68.1 | 72.9 | 77.1 | 82.6 | 87.2 | 93.6 |
| 50–64 | 33 | 79.4 | 8.2 | 62.5 | 71.1 | 75.0 | 80.8 | 85.4 | 93.5 |
| 65–74 | 38 | 76.2 | 7.5 | 64.6 | 68.9 | 70.8 | 75.8 | 83.3 | 89.6 |
| 75+* | 21 | 65.8 | 8.0 | 53.2 | 57.5 | 61.7 | 64.6 | 70.5 | 79.2 |
| **Tools** |  |  |  |  |  |  |  |  |  |
| 18–49 | 264 | 79.6 | 11.0 | 60.4 | 67.4 | 72.3 | 81.0 | 87.5 | 95.7 |
| 50–64 | 33 | 78.3 | 10.3 | 66.0 | 66.7 | 68.8 | 79.2 | 82.6 | 97.9 |
| 65–74 | 38 | 75.7 | 9.3 | 62.5 | 64.6 | 66.7 | 76.8 | 83.0 | 93.6 |
| 75+* | 21 | 72.0 | 10.0 | 54.2 | 62.5 | 63.8 | 72.9 | 78.7 | 85.4 |
| **Pseudowords** |  |  |  |  |  |  |  |  |  |
| 18–49 | 264 | 82.7 | 10.4 | 62.2 | 72.3 | 76.7 | 84.3 | 91.1 | 97.7 |
| 50–64 | 33 | 78.0 | 12.7 | 45.8 | 70.5 | 72.9 | 82.2 | 87.0 | 93.0 |
| 65–74 | 38 | 68.7 | 11.9 | 48.9 | 57.5 | 59.5 | 67.8 | 77.8 | 91.7 |
| 75+* | 21 | 64.0 | 11.6 | 50.0 | 53.3 | 56.2 | 63.0 | 68.9 | 80.8 |

Note. Values are percentage-correct scores collapsed across the Learning and Novel Stages. Cells in the 75+ age band (marked \*) should be interpreted with caution because *N* < 30.

Normative distributions generally shifted downward in older bands, especially for categories with stronger age associations. Equal raw percentages occupied different percentiles depending on category and age band, supporting interpretation of profiles against category-specific references. MoCA completers and non-completers showed no significant differences in the measured demographic or performance characteristics; the small administration-mode comparison was also nonsignificant. Supplementary Materials S3.3–S3.4 report estimates and CIs.

## Discussion

The ViRA-L extends a common recognition-memory structure across ten categories selected for neuropsychological and cortical relevance. Good battery-level reliability, broad score dispersion, and limited floor and ceiling effects support its use for normative characterization and individual-differences research. Lower accuracy and slower responses for novel views and difficult items provide converging evidence that performance reflects the intended recognition demands. Age was the clearest demographic correlate, but its association with accuracy differed markedly across categories, demonstrating the need for age-sensitive interpretation of individual recognition profiles.

The methodological contribution is the combination of category coverage and procedural comparability. Cambridge-style tests and the Vanderbilt Expertise Test established the value of matched formats (Dennett et al., 2012; Duchaine & Nakayama, 2006; McGugin et al., 2012). The ViRA-L extends this approach to domains implicated in neurological recognition disorders and neuroimaging research. Matching memory load, response format, and scored-trial structure reduces several confounds that otherwise complicate comparisons between faces, words, and objects. It therefore enables future patient studies to examine whether associations and dissociations persist under comparable demands. Category-level reliability was weaker than aggregate reliability, particularly for Cars, and should temper interpretation of isolated low scores.

The larger age association in Novel than Learning trials helps specify what the battery measures. Learning trials tested recently studied identities in relatively familiar views; Novel trials required those identities to be recognized across altered appearance. The latter therefore placed greater demands on representations that generalize beyond a studied image. Older adults can retain same-view discrimination while showing disproportionate difficulty matching faces or novel objects across viewpoints (Habak et al., 2008; Pilz et al., 2011). Providing moving views during learning can improve later face recognition across viewpoints, suggesting that representational breadth at encoding supports generalization (Maguinness & Newell, 2014). The larger later-life difference in Novel trials is consistent with this account, although the design does not separate encoding, retention, and transformation tolerance. These cross-sectional associations should not be interpreted as longitudinal rates of decline.

The category pattern likewise requires several possible explanations. Pseudowords and Hands showed the largest modeled differences between ages 50 and 85, whereas Houses showed the smallest. Younger-adult baseline performance did not account for this ordering, suggesting that later-life differences were not simply greatest for categories that were already hardest. One possibility is perceptual expertise: recognition contains both shared ability and domain-specific components associated with experience (McGugin et al., 2012). Generations may differ in familiarity with particular cars, tools, instruments, or orthographic forms because of educational, occupational, or cultural experience. The Age × Category interaction may consequently combine age-associated differences with cohort differences in individuation expertise; the present cross-sectional design cannot disentangle them.

Semantic or overlearned support offers a related explanation. Pseudowords lack lexical meaning and cannot benefit from stored word representations. Real-word recognition can remain relatively preserved across adulthood, whereas age costs become more apparent when tasks emphasize orthographic analysis or sublexical decisions (Cohen-Shikora & Balota, 2016; Froehlich et al., 2016; Payne & Federmeier, 2018; Rojas et al., 2024). This could contribute to the pronounced pseudoword association. However, homogeneous subcategories reduced reliance on broad semantic labels throughout the battery. Semantic support therefore cannot, by itself, explain why pseudowords differed from every other category, and future studies should manipulate semantic availability within a matched recognition task.

Developmental timing and representational distinctiveness provide another account. Category representations emerge on different schedules: infant evidence indicates early face representations, later differentiation of limb and character-like stimuli, and literacy-dependent specialization for words (Yan et al., 2024; Yeatman et al., 2024). Longitudinal childhood data further show increasing word and face distinctiveness alongside weakening limb representations, with distinctiveness predicting recognition (Nordt et al., 2023). Combined with delayed and less selective visual representations in older adults (Haupt et al., 2025), these findings motivate a last-in, first-out hypothesis: later or more experience-dependent specialization may be especially vulnerable (Douaud et al., 2014). Earlier establishment and lifelong reinforcement could contribute to the more moderate face association relative to limb-related and orthographic categories. Nevertheless, evidence for age-related dedifferentiation differs across tasks and category systems (Chee et al., 2006; Goh et al., 2010; Koen & Rugg, 2019; Srokova et al., 2020). Studies have reported both reductions in neural specificity and null effects, with differences in task demands and stimulus context complicating comparisons (D. C. Park et al., 2004; J. Park et al., 2012; Payer et al., 2006; Voss et al., 2008; Zebrowitz et al., 2016; Zheng et al., 2018). Our behavioral findings cannot establish this developmental mechanism.

Fine-grained perceptual overlap may also matter. Older adults have difficulty distinguishing similar exemplars and retrieving the perceptual detail needed to separate them (Boutet et al., 2015; Bowman et al., 2019; Burnside et al., 2017; Pidgeon & Morcom, 2014). The presemantic account of category effects similarly attributes some category differences to structural similarity and visual matching demands before semantic access (Bordaberry et al., 2016). Bodies and Hands may require particularly precise configural or part-based discrimination, whereas Houses may provide more stable global layout cues. We examined this prediction by relating average image similarity to category-specific age differences. Across nine image-based categories, the association was positive but nonsignificant (Supplementary Material S3.2). Pseudowords were excluded because their SSIM values did not provide a meaningful similarity measure. The result motivates further testing but does not confirm the account. Expertise, semantic support, developmental history, and perceptual overlap may act together rather than provide mutually exclusive explanations.

Gender and education require similarly cautious interpretation. Their nonsignificant overall accuracy coefficients do not imply that they are irrelevant across categories or populations. Only 25 participants had lower education, all in the older cohort; restricted variation limited sensitivity to education-related associations and interactions. An overall gender coefficient may also average over category-specific patterns. The Vanderbilt Expertise Test reported category-dependent sex effects (McGugin et al., 2012), a meta-analysis identified a modest female advantage and female own-gender bias in face recognition (Herlitz & Lovén, 2013), and education predicted visual long-term recognition memory in another normative study (Barvas et al., 2022). Age was the best-supported basis for the present norms, while demographic influences may differ with task and sample composition (delCacho-Tena et al., 2024).

The age-stratified norms provide empirical reference distributions for each category, whose scores are not interchangeable even before demographic adjustment. Such references can reduce overinterpretation of low scores in older adults and underrecognition of atypical performance in younger adults (Arampatzi et al., 2024; Karstens et al., 2024). They offer a foundation for research on posterior cerebral artery or occipitotemporal stroke, posterior cortical atrophy, prosopagnosia, alexia, and object agnosia. Maintaining age bands without further demographic subdivisions preserved reference-cell sizes and made their limitations visible. This choice reflects the available sample and the observed demographic associations, rather than a claim that gender and education never require adjustment (Brown et al., 2025; Karstens et al., 2024).

The small 75+ cell limits percentile stability, and ages 36–49 were unrepresented. Regarding the latter, the equal performance in the large young adults sample compared to the adults of 50 to 64 makes it unlikely that there would suddenly be a difference in the unsampled age range of 36-49. Nevertheless, while our test battery and norms are very useful for a wide range of sampling and grouping situations, particular care should be taken when a study would want to focus upon the range that we did not sample. The lower-education subsample and the oldest age band also address different questions: limited educational diversity constrains demographic adjustment, whereas a small normative cell constrains reference precision at those ages. Larger, more evenly distributed samples are needed to strengthen both aspects of the normative framework (O’Connell et al., 2022).

Administration and recruitment also constrain generalizability. Although remote cognitive testing can produce results comparable to in-person assessment (Backx et al., 2020; Latham et al., 2025; Requena-Komuro et al., 2022), only nine participants were tested on site, precluding a strong mode comparison (Supplementary Material S3.3). Physical display size, viewing distance, distractions, and breaks varied without objective monitoring. Online volunteers may have greater digital literacy, education, and independence than the wider older population or clinical samples. Incomplete MoCA screening adds uncertainty: neither the nonsignificant completer comparisons nor the stable age result after MoCA exclusions rules out unmeasured cognitive differences among unscreened participants.

Procedural matching also does not ensure visual equivalence. Categories were not equated on a global complexity measure and differed in shape, part structure, homogeneity, and meaningful transformations. Tools often contain elongated parts, whereas hands, faces, and bodies require discrimination of closely related configurations. Preserving these characteristics maintains the relevance of each category, but leaves residual stimulus variation that may contribute to performance. Category effects should therefore not be attributed exclusively to category-selective cognitive mechanisms. Native Dutch recruitment further limits cultural and linguistic generalization, especially for pseudowords and faces; using pseudowords and a broader face set than the CFMT mitigates but does not remove this concern. Expertise and familiarity can vary across cultures for other categories as well.

Several other follow-up studies are relevant. Direct comparison with established category-specific measures, category-level test–retest reliability, ecological validity, and measurement invariance across languages, devices, and administration modes remain to be established. The absence of a direct CFMT comparison is particularly relevant because shared task logic alone does not establish convergent validity. Administering the ten categories during functional imaging could test whether younger and older adults differ in neural differentiation across these domains (Koen & Rugg, 2019; Srokova et al., 2020). Patient studies are needed to establish known-groups validity, diagnostic accuracy, clinically meaningful cutoffs, and sensitivity to change. These extensions would connect the present normative characterization to the clinical interpretation of selective and generalized recognition difficulties.

## Conclusion

The ViRA-L combines ten neuropsychologically relevant categories within a matched recognition-memory framework and provides age-stratified normative references. Its reliability, score dispersion, and sensitivity to task demands support research use, while category-specific age associations show why recognition profiles require both demographic and domain-specific interpretation. The battery provides a basis for studying shared and selective recognition difficulties; comparison with established measures and validation in neurological populations will determine its clinical utility.

## Acknowledgements

We thank Brad Duchaine for advice and Silke Busschots, Fjolla Alijaj, and Amber Gordts for assistance with participant recruitment.

## Use of artificial intelligence

We used ChatGPT (OpenAI; GPT-5.6 Sol, accessed September-October 2026) for English-language editing, rephrasing, and clarification of author-written text. It was not used to generate research data or perform statistical analyses. The authors reviewed and revised all AI-assisted text and take full responsibility for the manuscript.

## Availability of materials

Task materials, analysis scripts, norms, and documentation will be available through HOPLAB GitHub. The complete stimulus set cannot be distributed openly because third-party materials remain subject to their original licenses. Permitted stimuli will be available for scientific use under those conditions; the repository will identify original sources and procedures for obtaining or reproducing restricted materials.

