## Supplementary materials for "The ViRA-L (Visual Recognition Assessment–Leuven): Development, Validation, and Age-Stratified Normative Characterization of a Multi-Category Visual Recognition Battery"

Contents: S1–S2, stimulus and trial construction; S3, additional analyses; S4, extended methods; S5, detailed results; Tables S1–S14; Figures S1–S2; supplementary references.

#### S1. Stimulus construction and standardization

The ViRA-L comprised Cars, Bodies, Musical Instruments, Pseudowords, Tools, Faces, Food, Houses, Hands, and Animals. Six target identities and a larger distractor pool were selected per category. Image-based stimuli came from established datasets and publicly available three-dimensional models, with subsequent editing and rendering for the task.

Cars were drawn primarily from the Cambridge Car Memory Test stimulus set (Dennett et al., 2012), supplemented with vehicle models from Sketchfab and edited in Autodesk 3ds Max (Autodesk, 2022). Bodies were generated with MakeHuman (MakeHuman Community, 2023). Face stimuli were derived from the Multi-Racial Mega-Resolution (MR2) face database (Strohminger et al., 2016), Hands from Hand-3D-Studio (Zhao et al., 2020), and Houses from Houses3K (Peralta et al., 2020). Musical Instruments, Tools, Food, and Animals used Sketchfab models (Sketchfab, 2026), edited and rendered in Autodesk 3ds Max.

Relatively homogeneous subcategories increased visual similarity and reduced the usefulness of coarse semantic labels. Cars comprised sedans, sports cars, and wagons; Bodies, male and female bodies; Musical Instruments, guitars, brass instruments, and flutes; Tools, hammers, wrenches, and screwdrivers; Faces, African, East Asian, and Western identities; Food, burgers, apples, and bread; Houses, two architectural geometries; Hands, open and closed postures; and Animals, cats and dogs. Stimulus selection, subcategory composition, and transformation parameters were refined during piloting to avoid pronounced floor or ceiling effects. This procedure did not equate categories on a global measure of visual complexity.

Viewpoint and illumination changes were produced in Autodesk 3ds Max. Faces, Cars, Bodies, Houses, Animals, Hands, and Musical Instruments primarily used leftward and rightward rotations from a canonical view. Tools and Food used front-to-back rotations where these better altered diagnostically relevant surfaces or part relations. Selecting transformations according to object geometry preserved meaningful recognition demands; an identical rotation axis would not have produced equivalent changes across categories. Lighting changes further reduced the usefulness of exact image matching. Viewpoint and illumination were not fully crossed, so the analyses characterize combined appearance conditions rather than their independent effects.

Pseudowords were generated separately with Wuggy (Keuleers & Brysbaert, 2010), using the Dutch templates “volks” and “boter.” Candidates matched template length, subsyllabic segment lengths, and transition frequencies and were required to match two of the three subsyllabic segments. Font changes replaced lighting changes, using “Fat Wandals,” “Tha Funk – Urban Hook – UPZ,” and “Helvair Graffiti,” obtained from FontSpace (Grebäck et al., 2026). This introduced appearance variation while retaining orthographically legal, nonlexical strings.

Images were resized without changing aspect ratio and centered on an 800 × 600-pixel canvas, using the smaller horizontal or vertical scaling factor to fit the frame. Grayscale conversion was followed by multiplicative brightening of images below a predefined mean-luminance threshold; brighter images were unchanged. This reduced extreme brightness differences while preserving internal luminance structure. The PsychoPy window used height units and image components used normalized window units (Peirce et al., 2019). Presentation size therefore scaled with the browser window rather than being fixed in centimeters or visual angle.

#### S2. Trial construction and difficulty manipulation

For image categories, grayscale stimulus pairs were compared within category using the Structural Similarity Index Measure (SSIM; Wang et al., 2004). Rank-ordered pairwise scores were divided into the upper and lower halves. Difficult trials used more similar target–distractor pairings and easy trials less similar pairings, with final assignments refined during piloting. Pseudoword difficulty was defined by orthographic similarity: difficult distractors more closely preserved the target's letter structure. SSIM was not used to assign pseudoword difficulty.

Each category contained 18 scored Learning trials and 30 scored Novel trials. Each target contributed three Learning and five Novel recognition trials. Study presentations and the six-target review were separate, unscored events. Each scored trial presented one target and two distractors, yielding 96 distractor presentations per category. Distractor identities recurred across trials, reducing the usefulness of familiarity alone, consistent with the Cambridge Face Memory Test (CFMT; Duchaine & Nakayama, 2006).

### S3. Additional analyses

#### S3.1. Empirical lower reaction-time cutoffs

Lower reaction-time thresholds were estimated separately for students and older adults using kernel-density estimates of log reaction time for correct and incorrect responses. The selected lower-tail crossover was 590.87 ms for students and 738.73 ms for older adults, reported as 591 and 739 ms. Figures S1–S2 show the cutoffs and excluded faster responses. A Sheather–Jones bandwidth check gave 590 and 730 ms, respectively, indicating limited sensitivity to the smoothing specification. These thresholds were applied during trial preprocessing; reaction time was not analyzed as an outcome in the category-order comparison.

**Supplementary Figure S1**

*Correct- and incorrect-response RT densities in students*

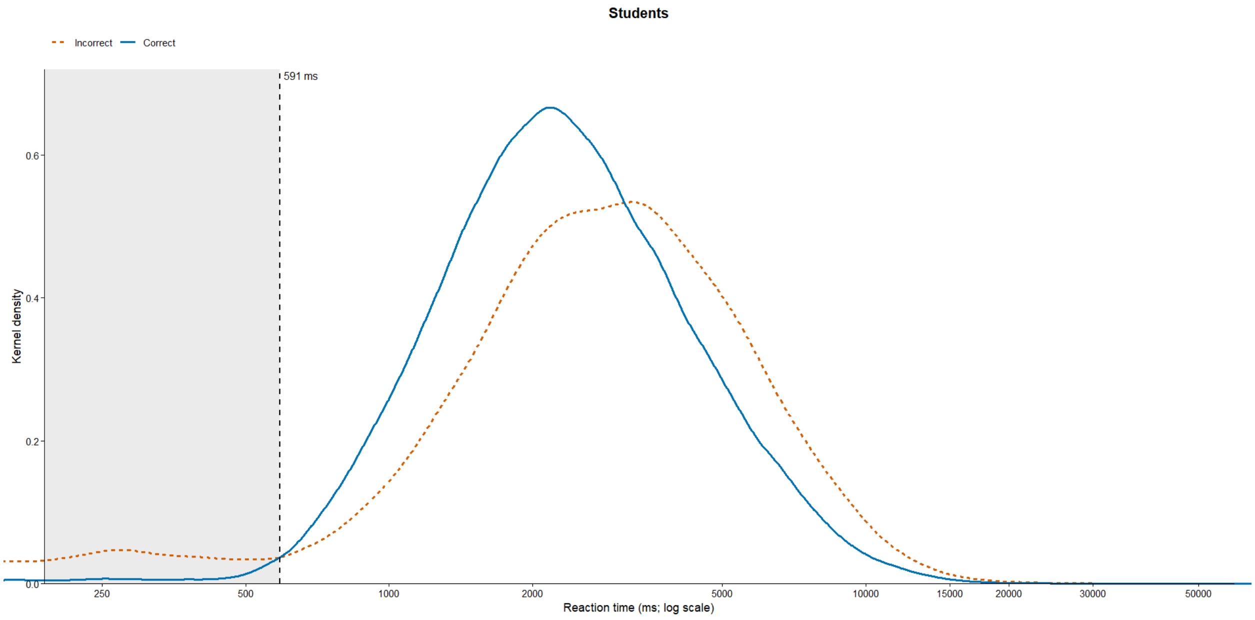


Note. The dashed line marks the selected 591-ms student cutoff; the shaded region contains excluded fast responses.

**Supplementary Figure S2**

*Correct- and incorrect-response RT densities in older adults*

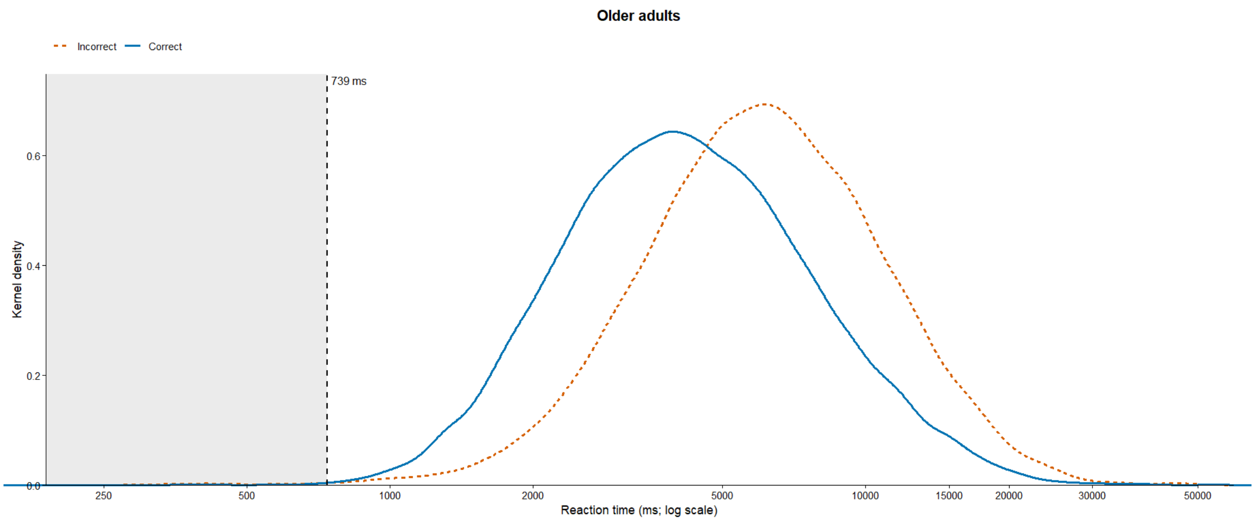


Note. The dashed line marks the selected 739-ms older-adult cutoff; the shaded region contains excluded fast responses.

**S3.2. Perceptual distinctiveness and later-life accuracy differences**

This exploratory analysis asked whether image categories with more visually distinctive exemplars showed smaller modeled later-life accuracy differences. The outcome was predicted accuracy at age 50 minus predicted accuracy at age 85, expressed in percentage points; larger positive values indicate a larger later-life difference. The same category estimates used in the main manuscript and Table S4 were used throughout this analysis. These are cross-sectional model predictions, not within-person changes.

Two category-level predictors were examined: SSIM-based distinctiveness, defined as one minus mean within-category SSIM (Wang et al., 2004), and mean DreamSim distance, a learned perceptual-distance measure (Fu et al., 2023). Higher values of either predictor indicate greater distinctiveness. DreamSim captures aspects of image appearance beyond pixel-level structural similarity, but neither measure should be treated as a direct measure of neural representational distinctiveness. The analysis used the category-level image summaries reported in Table S11. Pseudowords were excluded because their template-based construction makes their SSIM summary an unsuitable measure of the perceptual overlap relevant to this comparison. Both predictors were therefore compared over the same nine image categories.

Pearson correlations were accompanied by 95% Fisher-transformed CIs. Spearman rank correlations provided a robustness check, with average ranks for ties and two-sided asymptotic p values. The two distinctiveness measures were weakly related, $r(7) = .182$, 95% CI [−.548, .755], $p = .639$; Spearman $\rho = .159$, $p = .683$. SSIM-based distinctiveness was negatively associated with difference magnitude, $r(7) = −.521$, 95% CI [−.880, .219], $p = .150$; Spearman $\rho = −.383$, $p = .308$. Equivalently, mean SSIM similarity had a positive correlation of .521, 95% CI [−.219, .880]. DreamSim distance was also negatively associated with difference magnitude, $r(7) = −.505$, 95% CI [−.876, .239], $p = .165$; Spearman $\rho = −.527$, $p = .145$.

Separate ordinary least-squares regressions predicted the age-difference magnitude from each predictor, standardized using its sample mean and standard deviation across nine categories. A joint model included both standardized predictors. Slope coefficients therefore represent percentage-point differences per one-SD increase in a predictor; the outcome itself was not standardized. Coefficient CIs used the t distribution with seven residual degrees of freedom for single-predictor models and six for the joint model. All perceptual-analysis p values are exploratory and unadjusted.

The SSIM model yielded $b = −2.639$, 95% CI [−6.506, 1.227], $p = .150$, $R^2 = .271$, and adjusted $R^2 = .167$. The DreamSim model yielded $b = −2.560$, 95% CI [−6.469, 1.349], $p = .165$, $R^2 = .255$, and adjusted $R^2 = .149$. Leave-one-category-out prediction errors were similar: RMSE = 5.174 percentage points for SSIM and 5.282 for DreamSim. The joint model explained 44.5%

of between-category variance, adjusted $R^2 = .260$, $F(2, 6) = 2.41$, $p = .171$. Neither conditional slope was significant: SSIM, b = −2.248, 95% CI [−6.082, 1.587], $p = .201$; DreamSim, b = −2.150, 95% CI [−5.985, 1.684], $p = .219$. Table S12 gives all coefficients and model-comparison statistics.

AIC and BIC were computed from the Gaussian likelihood, counting the residual variance among estimated parameters. Residual SD uses residual degrees of freedom; leave-one-category-out RMSE is the square root of the mean squared prediction error over the nine held-out categories. These quantities have different denominators and are labeled separately. Small differences in fit and predictive error do not establish the superiority of either metric. With only nine categories, wide CIs, and uncertainty in the modeled age differences not propagated into these regressions, the results remain inconclusive. Their direction is consistent with the perceptual-overlap account, but they neither confirm that account nor exclude it.

**S3.3. On-site versus remote administration**

Administration mode was closely associated with cohort: students were tested remotely and all nine on-site participants were older adults. Comparisons were therefore restricted to the older cohort, with 86 online and nine on-site participants. Welch tests compared continuous outcomes; differences below are online minus on-site.

Online participants had a mean age of 67.84 years (*SD* = 8.46), compared with 71.11 (*SD* = 6.45) on site: difference = −3.27 years, 95% CI [−8.41, 1.87], $t(11.22) = -1.40$, $p = .190$. Age was available for 83 online participants and all nine on-site participants. Total accuracy was 74.82% (*SD* = 6.78) online and 73.86% (*SD* = 4.92) on site: difference = 0.96 percentage points, 95% CI [−2.97, 4.89], $t(11.46) = 0.53$, $p = .604$. Mean correct-response reaction time was 4.97 s

(*SD* = 1.43) online and 5.75 s (*SD* = 1.65) on site: difference = −0.78 s, 95% CI [−2.07, 0.51], $t(9.30) = -1.36$, $p = .205$.

These nonsignificant comparisons do not establish equivalence. The on-site sample was small, and the age difference between modes also remains imprecisely estimated. These reaction-time comparisons concern administration mode; they are separate from the accuracy-only category-order analysis.

**S3.4. MoCA completion and exclusion sensitivity**

Among the 95 retained older adults, 50 completed the MoCA and 45 did not. The completer group comprised 29 telephone MoCA-22 and 21 full-version administrations. Continuous variables were compared with Welch tests. Differences below are completers minus non-completers.

Mean age was 68.22 years (*SD* = 8.27) in completers and 68.10 (*SD* = 8.46) in non-completers: difference = 0.12 years, 95% CI [−3.36, 3.61], $t(86.57) = 0.07$, $p = .943$. Total accuracy was 74.55% (*SD* = 6.49) versus 74.93% (*SD* = 6.81): difference = −0.38 percentage points, 95% CI [−3.10, 2.33], $t(90.81) = -0.28$, $p = .780$. Mean correct-response reaction time was 5.05 s (*SD* = 1.39) versus 5.04 s (*SD* = 1.56): difference = 0.01 s, 95% CI [−0.59, 0.62], $t(88.69) = 0.04$, $p = .971$. Age, gender, and education were available for 42 of the 45 non-completers and all 50 completers; accuracy and reaction time were available for all 95.

The percentage male was 40.0% among completers and 38.1% among non-completers (20/50 versus 16/42; Fisher exact $p = 1.000$). Higher education was reported by 78.0% versus 66.7% (39/50 versus 28/42; $\chi^2(1) = 0.96$, $p = .326$). Unadjusted odds ratios and 95% Wald CIs are reported with these comparisons in Table S13. The results do not demonstrate equivalence or exclude unmeasured cognitive differences among unscreened participants.

The primary age model was re-estimated before and after MoCA-based exclusions. Before exclusions, the model included 358 participants and 167,158 trials, and the age likelihood-ratio test was $\chi^2(3) = 50.29$, $p < .001$. After exclusions, it included 355 participants and 165,735 trials, $\chi^2(3) = 49.39$, $p < .001$. The model-specific difference is three participants because one of the four MoCA-excluded participants was already absent from the covariate-complete model sample. The overall age association was therefore retained, although this sensitivity analysis cannot establish the cognitive status of unscreened participants.

**S3.5. Younger-adult accuracy and later-life differences**

Mean category accuracy among the 264 younger participants with known age was correlated with the category-specific age-50-minus-age-85 difference magnitude. Across ten categories, Pearson $r(8) = -.283$, 95% CI [−.775, .422], $p = .428$; Spearman $\rho = -.370$, $p = .296$. The negative estimate indicates that easier categories in younger adults tended to have smaller modeled later-life differences, but the wide CI includes substantial associations in either direction. This analysis does not establish that baseline difficulty explains, or fails to explain, the ordering of category-specific age differences.

#### S4. Extended methods and analytical specifications

##### S4.1. Administration and presentation

The task followed CFMT recognition-memory logic across all ten categories (Duchaine & Nakayama, 2006). Practice presented one target in three views and included three three-alternative forced-choice trials with feedback. For each of six study targets, three images were shown sequentially for 3,000 ms each, with 500-ms intervals, followed by three recognition trials. The resulting 18 Learning trials were followed by a 20-s, 2 × 3 display of the six targets and 30 Novel trials. Test displays remained until response; intertrial intervals were 500 ms. Number keys mapped to the top, middle, and bottom alternatives, and participants used their preferred hand. Only practice provided feedback.

Novel trials presented previously learned identities under changed orientation, illumination, or both. Older adults additionally received five motivational easy trials per category during this stage. These used frontal views and visually dissimilar distractors and were excluded from every analysis. Standard category scores therefore comprised at most 48 valid trials. Table S10 reports predictions for the coded transformation conditions included in the orientation models.

Presentation files were 300 × 225 or 250 × 188 pixels following stimulus preparation. PsychoPy image components were centered against dark gray and scaled relative to the window in normalized units (Peirce et al., 2019). Physical display dimensions and visual angle were not fixed. On-site sessions were supervised by an experimenter. Remote participants were instructed to use a computer in a quiet environment, maintain a comfortable viewing distance, and avoid interruptions. Everyone completed the battery within one day. Students completed one continuous task; older adults completed two parts with an optional break. Compliance, screen characteristics, viewing distance, distractions, and break duration were not objectively monitored. The order-control version randomized category blocks while retaining the same within-block trial order.

**S4.2. Preprocessing and sample accounting**

The main analyses were conducted in R 4.5.2 (R Core Team, 2025). From 217,222 raw rows, preprocessing removed one trial with missing accuracy or reaction time, 66 trials with nonpositive reaction times, 5,140 motivational easy trials, and 1,361 duplicate rows. Duplicates arose when participants repeated online task sections because they were uncertain that responses had been saved. Identification used participant, block, experiment part, trial number, and target, retaining the first occurrence.

Reaction times exceeding each participant's mean by three standard deviations removed 3,335 trials: 2,631 in students and 704 in older adults. The cohort-specific lower thresholds described in S3.1 then removed 6,405 trials: 6,324 across 202 students and 81 across nine older adults. This left 200,914 trials. The density approach was motivated by evidence on implausibly fast responses and response-time distributions (Crouzet et al., 2014; Voormann & Miller, 2024; Voskuilen et al., 2018).

Participant exclusions followed trial filtering. Sixteen participants retained fewer than 60% of trials in at least four categories; two additional participants performed below chance in more than one category. Four older adults scored below the ninth percentile on version-appropriate MoCA norms. Full MoCA scores used Dutch age-, sex-, and education-adjusted norms (Kessels et al., 2022); telephone MoCA-22 scores used demographically adjusted norms for that version (Ratcliffe et al., 2024).

The cleaned dataset contained 417 participants and 194,635 trials: 322 students and 95 older adults. Primary fixed-order analyses included 360 participants and 168,027 trials, comprising 265 students and 95 older adults. The 57 randomized-order students contributed only to the order comparison. The primary accuracy model included 355 participants with complete covariates and 165,735 trials; the normative sample required age but not all model covariates and included 356. Thus, psychometric, normative, age-model, and order-model sample sizes differ for specified reasons rather than reflecting additional undisclosed exclusions.

**S4.3. Scoring and psychometric definitions**

Category accuracy was the percentage of valid trials answered correctly within a category, combining Learning and Novel trials. Total accuracy was computed over all analyzable trials for each participant. Reaction-time summaries and models used correct responses; reaction times were log-transformed for mixed modeling.

Cronbach's α, omega total, and Spearman–Brown-corrected odd–even split-half reliability were computed across the ten category scores for the battery and six target-item scores within each category. Cronbach's α was accompanied by Feldt 95% CIs (Table S1): the battery interval was [.860, .897], and category intervals extended from [.376, .548] for Cars to [.714, .793] for Pseudowords. An item refers to a target identity represented across multiple trials, not an individual recognition trial. Item performance was the mean item score across participants, with higher values indicating easier items. Each corrected item–total correlation related an item to the mean of the other items in the same category. Table S2 summarizes item means, dispersion, and corrected item–total correlations.

Total and category score distributions were summarized by N, mean, standard deviation, minimum, maximum, skewness, and Shapiro–Wilk p values. Ceiling scores were 100% correct and floor scores were at or below the three-alternative forced-choice chance level of 33.3%. Table S3 includes counts and percentages to distinguish rare extremes from broad score clustering. Item-score averaging and valid-trial-weighted category accuracy are distinct summaries and need not have identical means.

**S4.4. Models, effect sizes, and confidence intervals**

Trial accuracy was modeled with a binomial distribution and logit link, including age, gender, education, stage, category, and difficulty as fixed effects and participant and item random intercepts. Correct-trial log reaction time was modeled with corresponding linear mixed-effects models. Reference levels were female gender, lower education, Learning Stage, Cars, and easy difficulty. Tables S7–S8 report fixed-effect estimates, standard errors, confidence intervals, and tests. Age spline-basis coefficients describe a joint nonlinear function and should not be interpreted individually as effects per year.

For accuracy, linear age, three-degree, and four-degree natural spline models had AIC values of 147,733.59, 147,714.86, and 147,716.98, respectively. The three-degree spline was selected. For reaction time, corresponding AICs were 162,949.58, 162,949.82, and 162,949.18: all were within one AIC unit. The three-degree reaction-time model retained the same age specification for consistency; it was not uniquely favored by AIC.

Likelihood-ratio comparisons tested overall age effects and interactions. The significant accuracy Age × Category interaction was followed by category-specific models; all ten category-specific age tests survived false-discovery-rate correction. Predicted accuracies and age-85-versus-age-50 odds ratios summarized the nonlinear associations (Table S4). Age × Stage and Age × Difficulty were evaluated for accuracy and reaction time; Age × Education was additionally assessed for accuracy. Stage × Category tested heterogeneity in Learning–Novel differences, followed by estimated marginal contrasts within category with Holm correction across ten p values (Table S9). Table S5 collects the omnibus tests. Table S14 gives all pairwise orientation contrasts, including their odds ratios and multiplicity-adjusted intervals.

Accuracy coefficient CIs were calculated as $\beta \pm 1.96 \times SE$, and odds-ratio CIs by exponentiating these limits. Reaction-time fixed-effect CIs also used the normal Wald critical value, matching the supplied effect-size export; the t tests retain coefficient-specific Satterthwaite degrees of freedom. Estimated marginal age contrasts for reaction time use asymptotic inference. Exponentiation gives geometric-mean reaction-time ratios and corresponding intervals. Stage and age contrasts use the intervals exported with the estimated marginal contrasts. Age and Learning–Novel contrast intervals are pointwise 95% CIs. Orientation pairwise contrasts use Holm-adjusted p values with Bonferroni-adjusted 95% CIs within each stage (three Learning and ten Novel contrasts). Because Holm and Bonferroni are different adjustments, an interval may include the null even when its Holm-adjusted p value is below .05. Category-specific order contrasts use Holm-adjusted p values and pointwise CIs. False-discovery-rate corrections apply to the specified omnibus p-value families.

Predicted accuracies at individual ages or transformation conditions have their own pointwise CIs (Table S10). These are not CIs for differences between predictions. Percentage-point contrasts and their CIs were obtained by regridding the estimated marginal predictions to the response scale before calculating differences, thereby retaining the modeled covariance. The supplied contrast output provides overall, category-specific, and stage-specific age differences and the stage difference of differences (Tables S4B and S10D). Intervals were not reconstructed by subtracting separate prediction-CI endpoints. Omnibus likelihood-ratio statistics are accompanied by model-based estimates and contrasts rather than being relabeled as standardized effect sizes.

Orientation effects were evaluated separately within Learning and Novel stages by comparing models with and without the coded orientation condition. Within-category subcategory tests used logistic mixed models with participant and item random intercepts and difficulty and orientation as covariates; false-discovery-rate adjustment covered ten omnibus tests (Table S6). The subcategory lookup matched all 168,027 fixed-order trial rows.

Participant-level speed–accuracy analyses correlated total accuracy with mean correct-response reaction time and its within-participant standard deviation, using Pearson coefficients and Fisher-transformed 95% CIs. Welch comparisons used raw mean differences with t-based CIs. Categorical MoCA comparisons retained Fisher or $\chi^2$ tests from the original analyses; associated descriptive odds-ratio CIs were computed from the $2 \times 2$ cell counts on the log-odds scale. The exploratory perceptual regressions and interval calculations were independently recomputed in Python from the supplied category summaries and full-precision estimates; the original fitted mixed-model results were retained.

##### S4.5. Normative framework

Norms used first-session fixed-order data after trial and participant exclusions. Four participants without age information, including three older adults and one student, were omitted, leaving $N = 356$. Scores combined Learning and Novel trials within each category. No total-battery normative table was included because the intended interpretation concerns category profiles.

Prespecified age bands were 18–49, 50–64, 65–74, and 75+. Age was the clearest overall demographic correlate of accuracy. Further subdivision by gender and education would have reduced reference-cell stability and was not adopted for the principal tables. Main Table 2 reports N, mean, standard deviation, and the 5th, 16th, 25th, 50th, 75th, and 95th percentiles in each category-by-age-band cell, flagging $N < 30$.

Grouped empirical norms make the supporting sample in each cell visible and avoid continuous regression-based interpolation through an age interval without observations. Participants assigned to the nominal 18–49 band were actually aged 18–35. These norms do not directly establish expected performance for ages 36–49, and the nonsignificant comparison with ages 50–64 cannot fill that gap.

#### S5. Detailed results

##### S5.1. Score distributions and item functioning

Total accuracy in the fixed-order sample was $M$ = 78.69%, $SD$ = 7.01, range = 54.58%–95.91%, with skewness −0.48. Category means ranged from 65.22% for Cars to 86.68% for Animals (Table S3). Three participants scored at or below chance for Animals and one each for Cars, Bodies, Pseudowords, and Hands. Perfect scores occurred for nine participants in Animals, five in Houses, three in Tools, two each in Musical Instruments, Pseudowords, and Hands, and one in Bodies. No perfect scores occurred for Cars, Faces, Food, or the total battery.

The item-level review flagged three Cars targets, one Bodies target, and one Musical Instruments target. Flags concerned low performance or low corrected item–total correlations and were retained as indicators for future refinement, not used to remove additional items from this analysis. Mean corrected item–total correlations were lowest for Cars (.253) and highest for Pseudowords (.525). Good aggregate reliability does not remove the need to improve and validate individual category scales.

##### S5.2. Task and demographic coefficients

In the main accuracy model, Novel versus Learning yielded $\beta = -1.779$, $SE = 0.018$, $z = -97.55$, $OR = 0.169$, 95% CI [0.163, 0.175], $p < .001$. Difficult versus easy yielded $\beta = -0.561$, $SE = 0.014$, $z = -39.70$, $OR = 0.571$, 95% CI [0.555, 0.587], $p < .001$. Gender yielded $\beta = 0.007$, $SE = 0.063$, $z = 0.12$, $OR = 1.008$, 95% CI [0.890, 1.141], $p = .906$; education yielded $\beta = 0.149$, $SE = 0.102$, $z = 1.46$, $OR = 1.160$, 95% CI [0.950, 1.418], $p = .145$. Full coefficients are in Table S7.

For log reaction time, Novel versus Learning yielded $\beta = 0.463$, $SE = 0.003$, 95% CI [0.457, 0.469], $t(123{,}994.63) = 152.18$, $p < .001$, corresponding to a geometric-mean ratio of

1.589, 95% CI [1.579, 1.598]. Difficult versus easy yielded $\beta = 0.161$, *SE* = 0.003, 95% CI [0.156, 0.166], $t(130{,}032.08) = 61.64$, $p < .001$; ratio = 1.175, 95% CI [1.169, 1.181]. Higher education yielded $\beta = 0.142$, *SE* = 0.052, 95% CI [0.040, 0.243], $t(354.89) = 2.73$, $p = .007$; ratio = 1.152, 95% CI [1.041, 1.275]. Gender yielded $\beta = -0.026$, *SE* = 0.032, 95% CI [−0.088, 0.036], $t(354.60) = -0.82$, $p = .413$; ratio = 0.974, 95% CI [0.915, 1.037]. Table S8 reports the full model. The overall age-85-versus-age-50 geometric-mean reaction-time ratio was 1.43, 95% CI [1.19, 1.71], $z = 3.864$, $p < .001$. Predicted geometric means were 3.33 s at age 50, 95% CI [3.01, 3.68], and 4.75 s at age 85, 95% CI [4.24, 5.32].

Total accuracy showed little association with mean correct-response reaction time, $r(358) = -.042$, 95% CI [−.145, .062], $p = .428$, or within-participant reaction-time variability, $r(358) = .054$, 95% CI [−.050, .156], $p = .308$. These analyses did not support a simple participant-level speed–accuracy trade-off.

The overall accuracy age contrast was *OR* = 0.370, 95% CI [0.260, 0.526], $z = -5.538$, $p < .001$, or −14.5 percentage points, 95% CI [−19.9, −9.0], $z = -5.192$, $p < .001$. Overall predicted accuracy was 88.6% at age 50, 95% CI [86.2, 90.5], and 74.1% at age 85, 95% CI [69.3, 78.4]. These estimated marginal predictions were averaged over gender, education, stage, category, and difficulty. Category-specific differences and CIs are given on both odds-ratio and percentage-point scales in Tables S4A–S4B; their p values differ because they test contrasts on different scales.

Learning accuracy predictions were 95.78%–96.96% across frontal and 30° views. Novel predictions ranged from 74.79% for rotation with lateral shadow to 81.58% for one frontal-shadow condition (Table S10A). Predicted age-50 and age-85 accuracy was 95.39% and 87.50% in Learning, a difference of −7.89 percentage points, and 75.70% and 53.91% in Novel, a difference of −21.79 (Table S10C). The corresponding age-difference CIs were [−11.1, −4.63] and [−29.6, −13.93] percentage points, respectively, both $p < .001$. The Novel-minus-Learning difference of age differences was −13.9 percentage points, 95% CI [−19.4, −8.36], $z = −4.92$, $p < .001$. On the odds scale, the ratio of the two age odds ratios was 1.11, 95% CI [0.886, 1.39], $z = 0.91$, $p = .365$. The significant omnibus Age × Stage test evaluates the age functions jointly; this nonsignificant odds-scale endpoint contrast asks a different question. The absolute probability-scale contrast can also be significant when its odds-scale counterpart is not, because interaction depends on scale.

**S5.3. Normative age-band comparison**

The normative sample included 264 participants aged 18–35 assigned to the 18–49 band, 33 aged 50–64, 38 aged 65–74, and 21 aged 75+. Mean total accuracy was 80.09% (*SD* = 6.60) in the youngest band and 78.70% (*SD* = 5.49) at ages 50–64. The youngest-minus-50–64 difference was 1.39 percentage points, 95% CI [−0.70, 3.48], $t(44.44) = 1.34$, $p = .188$. The standardized difference was Hedges' g = 0.21, 95% CI [−0.15, 0.57], using the pooled standard deviation. This comparison neither establishes equivalence nor validates extrapolation to the unobserved ages 36–49. The small 75+ group separately limits precision at the oldest ages.

**Supplementary Table S1**

*Battery- and category-level reliability*

| Scale / category | N | k | α | 95% CI for α | Omega total | Split-half |
|---|---|---|---|---|---|---|
| Battery | 360 | 10 | 0.879 | [0.860, 0.897] | 0.901 | 0.887 |
| Animals | 360 | 6 | 0.607 | [0.541, 0.667] | 0.762 | 0.657 |
| Bodies | 360 | 6 | 0.539 | [0.461, 0.609] | 0.778 | 0.556 |
| Cars | 360 | 6 | 0.466 | [0.376, 0.548] | 0.674 | 0.480 |
| Faces | 360 | 6 | 0.500 | [0.415, 0.576] | 0.687 | 0.596 |
| Food | 360 | 6 | 0.550 | [0.474, 0.619] | 0.740 | 0.610 |
| Hands | 360 | 6 | 0.617 | [0.552, 0.675] | 0.751 | 0.644 |
| Houses | 360 | 6 | 0.583 | [0.512, 0.646] | 0.753 | 0.619 |
| Musical Instruments | 360 | 6 | 0.511 | [0.429, 0.586] | 0.700 | 0.504 |
| Pseudowords | 360 | 6 | 0.756 | [0.714, 0.793] | 0.823 | 0.793 |
| Tools | 360 | 6 | 0.637 | [0.576, 0.692] | 0.729 | 0.660 |

Note. N denotes participants and k the number of category scales (battery) or target items (categories). Alpha CIs use the Feldt method. Split-half estimates are Spearman–Brown-corrected odd–even correlations.

**Supplementary Table S2**

*Category-level item statistics*

| Category | Items | Mean item score | SD | Min | Max | Mean corrected item–total r |
|---|---|---|---|---|---|---|
| Animals | 6 | 0.847 | 0.067 | 0.763 | 0.919 | 0.411 |
| Bodies | 6 | 0.798 | 0.054 | 0.720 | 0.849 | 0.316 |
| Cars | 6 | 0.604 | 0.138 | 0.445 | 0.730 | 0.253 |
| Faces | 6 | 0.699 | 0.063 | 0.611 | 0.786 | 0.279 |
| Food | 6 | 0.805 | 0.073 | 0.703 | 0.910 | 0.311 |
| Hands | 6 | 0.771 | 0.071 | 0.660 | 0.839 | 0.379 |
| Houses | 6 | 0.831 | 0.061 | 0.746 | 0.911 | 0.362 |
| Musical Instruments | 6 | 0.779 | 0.085 | 0.667 | 0.885 | 0.292 |
| Pseudowords | 6 | 0.788 | 0.047 | 0.702 | 0.836 | 0.525 |
| Tools | 6 | 0.774 | 0.068 | 0.666 | 0.844 | 0.395 |

Note. Higher item performance indicates an easier item. Corrected item–total correlations relate each target-item score to the mean of the remaining target items in the same category. Item-score means differ from valid-trial-weighted category accuracy.

**Supplementary Table S3**

*Score distributions and floor and ceiling counts*

| Scale / category | N | Mean | SD | Min | Max | Skew | Shapiro–Wilk p | Ceiling *n* (%) | Floor *n* (%) |
|---|---|---|---|---|---|---|---|---|---|
| Total score | 360 | 78.69 | 7.01 | 54.58 | 95.91 | −0.48 | < .001 | 0 (0.00) | 0 (0.00) |
| Animals | 360 | 86.68 | 9.62 | 16.67 | 100.00 | −2.81 | < .001 | 9 (2.50) | 3 (0.83) |
| Bodies | 360 | 79.91 | 9.53 | 31.11 | 100.00 | −0.86 | < .001 | 1 (0.28) | 1 (0.28) |
| Cars | 360 | 65.22 | 11.55 | 31.25 | 97.78 | 0.09 | .872 | 0 (0.00) | 1 (0.28) |
| Faces | 360 | 71.99 | 10.00 | 39.58 | 95.83 | −0.27 | .018 | 0 (0.00) | 0 (0.00) |
| Food | 360 | 80.65 | 7.86 | 45.65 | 97.92 | −0.76 | < .001 | 0 (0.00) | 0 (0.00) |
| Hands | 360 | 79.03 | 11.19 | 26.67 | 100.00 | −1.03 | < .001 | 2 (0.56) | 1 (0.28) |
| Houses | 360 | 84.69 | 9.23 | 43.59 | 100.00 | −1.17 | < .001 | 5 (1.39) | 0 (0.00) |
| Musical Instruments | 360 | 80.07 | 8.91 | 52.08 | 100.00 | −0.41 | < .001 | 2 (0.56) | 0 (0.00) |
| Pseudowords | 360 | 79.50 | 12.57 | 32.00 | 100.00 | −0.71 | < .001 | 2 (0.56) | 1 (0.28) |
| Tools | 360 | 78.57 | 10.89 | 43.75 | 100.00 | −0.32 | < .001 | 3 (0.83) | 0 (0.00) |

Note. Accuracy values are percentages. Ceiling denotes 100% accuracy and floor denotes accuracy at or below chance (33.3%).

**Supplementary Table S4A**

*Category-specific age estimates and odds-ratio contrasts*

| Category | Age 50 (%) | Age 85 (%) | Difference (pp) | OR | 95% CI for OR | p |
|---|---|---|---|---|---|---|
| Pseudowords | 88.35 | 65.09 | −23.26 | 0.246 | [0.158, 0.383] | < .001 |
| Hands | 89.48 | 69.03 | −20.45 | 0.262 | [0.168, 0.410] | < .001 |
| Cars | 77.02 | 59.18 | −17.84 | 0.433 | [0.284, 0.659] | < .001 |
| Bodies | 87.81 | 70.02 | −17.78 | 0.324 | [0.208, 0.505] | < .001 |
| Musical Instruments | 89.08 | 71.49 | −17.60 | 0.307 | [0.196, 0.482] | < .001 |
| Faces | 80.90 | 66.38 | −14.52 | 0.466 | [0.302, 0.719] | < .001 |
| Tools | 88.32 | 78.33 | −9.99 | 0.478 | [0.305, 0.749] | .001 |
| Food | 89.99 | 80.72 | −9.27 | 0.466 | [0.295, 0.734] | < .001 |
| Animals | 94.62 | 86.18 | −8.45 | 0.354 | [0.217, 0.580] | < .001 |
| Houses | 93.31 | 86.63 | −6.68 | 0.465 | [0.287, 0.752] | .002 |

Note. Differences are age 85 minus age 50 (percentage points). CIs and p values in this panel refer to odds ratios. Response-scale contrast CIs appear in S4B and prediction CIs in S10B. All intervals and tests are pointwise.

**Supplementary Table S4B**

*Category-specific age-85-minus-age-50 differences on the accuracy scale*

| **Category** | **Difference (pp)** | ***SE* (pp)** | **95% *CI* (pp)** | **z** | **p** |
|---|---|---|---|---|---|
| Pseudowords | −23.26 | 4.38 | [−31.80, −14.69] | −5.32 | < .001 |
| Hands | −20.45 | 4.14 | [−28.60, −12.33] | −4.93 | < .001 |
| Cars | −17.84 | 4.69 | [−27.00, −8.66] | −3.81 | < .001 |
| Bodies | −17.78 | 4.07 | [−25.80, −9.81] | −4.37 | < .001 |
| Musical Instruments | −17.60 | 3.98 | [−25.40, −9.79] | −4.42 | < .001 |
| Faces | −14.52 | 4.41 | [−23.20, −5.88] | −3.29 | < .001 |
| Tools | −9.99 | 3.39 | [−16.60, −3.36] | −2.95 | .003 |
| Food | −9.27 | 3.12 | [−15.40, −3.16] | −2.97 | .003 |
| Animals | −8.45 | 2.49 | [−13.30, −3.57] | −3.40 | < .001 |
| Houses | −6.68 | 2.43 | [−11.40, −1.92] | −2.75 | .006 |

Note. Differences and confidence limits are in percentage points. Values retain the precision available in the supplied response-scale contrast export; lower limits were exported to one decimal place. These CIs are pointwise, and p values are unadjusted. The p values differ from Table S4A because contrasts are tested on different scales.

**Supplementary Table S5**

*Omnibus likelihood-ratio tests*

| Outcome | Test | $\chi^2$ | df | p |
|---|---|---|---|---|
| Accuracy | Age overall | 49.39 | 3 | < .001 |
| Accuracy | Age × Category | 344.47 | 27 | < .001 |
| Accuracy | Age × Stage | 8.78 | 3 | .032 |
| Accuracy | Age × Difficulty | 1.17 | 3 | .760 |
| Accuracy | Age × Education | 1.15 | 2 | .564 |
| Accuracy | Stage × Category | 818.26 | 9 | < .001 |
| Accuracy, students | Order × Category | 33.14 | 9 | < .001 |
| Log reaction time | Age overall | 285.79 | 3 | < .001 |
| Log reaction time | Age × Stage | 345.45 | 3 | < .001 |
| Log reaction time | Age × Difficulty | 28.38 | 3 | < .001 |
| Accuracy | Orientation: Learning | 98.99 | 2 | < .001 |
| Accuracy | Orientation: Novel | 346.86 | 4 | < .001 |

Note. Tests compare nested models. Reaction time was not analyzed for order effects. Age × Education adds two estimable parameters in this sample; its reported degrees of freedom are retained from the fitted-model comparison.

**Supplementary Table S6**

*Within-category subcategory effects*

| Category | Subcategories | $\chi^2$ | df | p | FDR p |
|---|---|---|---|---|---|
| Bodies | 2 | 4.99 | 1 | .026 | .127 |
| Pseudowords | 2 | 4.61 | 1 | .032 | .127 |
| Food | 3 | 6.53 | 2 | .038 | .127 |
| Tools | 3 | 5.82 | 2 | .055 | .136 |
| Musical Instruments | 3 | 4.06 | 2 | .131 | .262 |
| Hands | 2 | 0.74 | 1 | .388 | .539 |
| Cars | 3 | 1.72 | 2 | .423 | .539 |
| Faces | 3 | 1.68 | 2 | .432 | .539 |
| Houses | 2 | 0.17 | 1 | .678 | .749 |
| Animals | 2 | 0.10 | 1 | .749 | .749 |

Note. Models included participant and item random intercepts and difficulty and orientation covariates. False-discovery-rate adjustment covered ten tests. No subcategory effect survived correction. Subcategory definitions appear in S1–S2.

**Supplementary Table S7**

*Fixed effects of the primary accuracy model*

| **Term** | **β** | **SE** | **95% CI for β** | **z** | **p** |
|---|---|---|---|---|---|
| Intercept | 2.049 | 0.176 | [1.703, 2.394] | 11.62 | < .001 |
| Age spline basis 1 | 0.638 | 0.202 | [0.242, 1.033] | 3.16 | .002 |
| Age spline basis 2 | 0.027 | 0.097 | [−0.164, 0.217] | 0.27 | .784 |
| Age spline basis 3 | −0.866 | 0.121 | [−1.103, −0.628] | −7.14 | < .001 |
| Male vs female | 0.007 | 0.063 | [−0.117, 0.132] | 0.12 | .906 |
| Higher vs lower education | 0.149 | 0.102 | [−0.051, 0.349] | 1.46 | .145 |
| Novel vs Learning | −1.779 | 0.018 | [−1.815, −1.744] | −97.55 | < .001 |
| Bodies vs Cars | 1.028 | 0.210 | [0.617, 1.438] | 4.91 | < .001 |
| Musical Instruments vs Cars | 0.940 | 0.210 | [0.528, 1.352] | 4.47 | < .001 |
| Pseudowords vs Cars | 0.954 | 0.211 | [0.540, 1.367] | 4.52 | < .001 |
| Tools vs Cars | 0.934 | 0.209 | [0.525, 1.344] | 4.48 | < .001 |
| Faces vs Cars | 0.443 | 0.209 | [0.033, 0.854] | 2.12 | .034 |

| **Term** | **β** | **SE** | **95% CI for β** | **z** | **p** |
|---|---|---|---|---|---|
| Food vs Cars | 1.113 | 0.208 | [0.705, 1.521] | 5.34 | < .001 |
| Houses vs Cars | 1.328 | 0.209 | [0.918, 1.737] | 6.36 | < .001 |
| Hands vs Cars | 0.872 | 0.211 | [0.459, 1.285] | 4.14 | < .001 |
| Animals vs Cars | 1.498 | 0.210 | [1.086, 1.909] | 7.13 | < .001 |
| Difficult vs easy | −0.561 | 0.014 | [−0.589, −0.533] | −39.70 | < .001 |

Note. $N$ = 355 participants; 165,735 trials. β is on the log-odds scale. Wald CIs use the normal critical value. Odds ratios and their CIs are obtained by exponentiating coefficients and limits. Reference levels are female, lower education, Learning, Cars, and easy difficulty. Age is a three-degree natural cubic spline, and its basis terms are interpreted jointly.

**Supplementary Table S8**

*Fixed effects of the correct-trial log-reaction-time model*

| Term | β | SE | 95% CI for β | df | t | p |
|---|---|---|---|---|---|---|
| Intercept | 0.663 | 0.068 | [0.530, 0.796] | 291.39 | 9.76 | < .001 |
| Age spline basis 1 | 0.391 | 0.103 | [0.188, 0.594] | 354.75 | 3.78 | < .001 |
| Age spline basis 2 | 0.672 | 0.049 | [0.576, 0.767] | 354.37 | 13.77 | < .001 |
| Age spline basis 3 | 0.815 | 0.062 | [0.694, 0.936] | 355.11 | 13.21 | < .001 |
| Male vs female | −0.026 | 0.032 | [−0.088, 0.036] | 354.60 | −0.82 | .413 |
| Higher vs lower education | 0.142 | 0.052 | [0.040, 0.243] | 354.89 | 2.73 | .007 |
| Novel vs Learning | 0.463 | 0.003 | [0.457, 0.469] | 123994.63 | 152.18 | < .001 |
| Bodies vs Cars | −0.213 | 0.059 | [−0.328, −0.098] | 59.94 | −3.63 | < .001 |
| Musical Instruments vs Cars | −0.256 | 0.059 | [−0.371, −0.140] | 59.98 | −4.35 | < .001 |
| Pseudowords vs Cars | −0.102 | 0.059 | [−0.217, 0.013] | 59.97 | −1.73 | .088 |
| Tools vs Cars | −0.304 | 0.059 | [−0.419, −0.189] | 59.97 | −5.18 | < .001 |
| Faces vs Cars | −0.370 | 0.059 | [−0.485, −0.255] | 60.05 | −6.29 | < .001 |

| **Term** | **β** | **SE** | **95% CI for β** | **df** | **t** | **p** |
|---|---|---|---|---|---|---|
| Food vs Cars | −0.620 | 0.059 | [−0.735, −0.505] | 59.93 | −10.55 | < .001 |
| Houses vs Cars | −0.430 | 0.059 | [−0.545, −0.315] | 59.92 | −7.32 | < .001 |
| Hands vs Cars | −0.229 | 0.059 | [−0.344, −0.114] | 60.00 | −3.90 | < .001 |
| Animals vs Cars | −0.589 | 0.059 | [−0.704, −0.474] | 59.92 | −10.03 | < .001 |
| Difficult vs easy | 0.161 | 0.003 | [0.156, 0.166] | 130032.08 | 61.64 | < .001 |

Note. CIs are normal Wald intervals, matching the effect-size export. The t tests use coefficient-specific Satterthwaite degrees of freedom. Exponentiating β and its limits gives geometric-mean reaction-time ratios and CIs. Reference levels match Table S7. The higher-education standard error is 0.05178056, rounded here to 0.052.

**Supplementary Table S9**

*Novel-versus-Learning accuracy contrasts within category*

| Category | β | SE | OR | 95% CI for OR | z | Holm p |
|---|---|---|---|---|---|---|
| Cars | −1.222 | 0.045 | 0.295 | [0.270, 0.322] | −27.24 | < .001 |
| Bodies | −2.123 | 0.059 | 0.120 | [0.107, 0.134] | −36.13 | < .001 |
| Musical Instruments | −1.640 | 0.059 | 0.194 | [0.173, 0.218] | −27.79 | < .001 |
| Pseudowords | −2.470 | 0.064 | 0.085 | [0.075, 0.096] | −38.31 | < .001 |
| Tools | −1.192 | 0.051 | 0.304 | [0.275, 0.335] | −23.37 | < .001 |
| Faces | −2.268 | 0.054 | 0.103 | [0.093, 0.115] | −42.14 | < .001 |
| Food | −2.687 | 0.070 | 0.068 | [0.059, 0.078] | −38.44 | < .001 |
| Houses | −1.385 | 0.063 | 0.250 | [0.221, 0.283] | −22.04 | < .001 |
| Hands | −1.358 | 0.054 | 0.257 | [0.231, 0.286] | −25.02 | < .001 |
| Animals | −1.949 | 0.088 | 0.142 | [0.120, 0.169] | −22.14 | < .001 |

Note. *OR* < 1 indicates lower Novel accuracy. CIs are pointwise 95% intervals; p values are Holm-adjusted across ten category contrasts. Every adjusted p value is below .001.

**Supplementary Table S10A**

*Predicted accuracy by stage and transformation condition*

| Stage | Condition | Predicted accuracy (%) | 95% CI |
|---|---|---|---|
| Learning | Frontal | 95.78 | [94.28, 96.90] |
| Learning | Left30 | 96.96 | [95.86, 97.78] |
| Learning | Right30 | 96.92 | [95.80, 97.74] |
| Novel | FrontalS1 | 81.58 | [78.13, 84.59] |
| Novel | FrontalS2 | 80.90 | [77.37, 84.00] |
| Novel | Left60 | 77.52 | [73.57, 81.04] |
| Novel | Right60 | 76.37 | [72.29, 80.02] |
| Novel | RightS1 | 74.79 | [70.55, 78.62] |

Note. The condition codes are preserved from the model export. Left30/Right30 denote 30° rotations; Left60/Right60 denote 60° rotations. FrontalS1/FrontalS2 are frontal-shadow conditions and RightS1 is a rotated-shadow condition. These are pointwise CIs for predicted means, not pairwise contrasts.

**Supplementary Table S10B**

*Pointwise accuracy predictions at ages 50 and 85 by category*

| Category | Age 50 accuracy (%) | 95% CI | Age 85 accuracy (%) | 95% CI |
|---|---|---|---|---|
| Pseudowords | 88.35 | [83.76, 91.78] | 65.09 | [55.39, 73.68] |
| Hands | 89.48 | [85.25, 92.60] | 69.03 | [59.71, 77.01] |
| Cars | 77.02 | [69.95, 82.83] | 59.18 | [49.53, 68.16] |
| Bodies | 87.81 | [83.07, 91.36] | 70.02 | [60.84, 77.84] |
| Musical Instruments | 89.08 | [84.71, 92.32] | 71.49 | [62.48, 79.06] |
| Faces | 80.90 | [74.32, 86.11] | 66.38 | [56.82, 74.75] |
| Tools | 88.32 | [83.71, 91.76] | 78.33 | [70.57, 84.50] |
| Food | 89.99 | [85.92, 92.98] | 80.72 | [73.50, 86.33] |
| Animals | 94.62 | [92.16, 96.34] | 86.18 | [80.30, 90.51] |
| Houses | 93.31 | [90.37, 95.40] | 86.63 | [80.91, 90.83] |

Note. Intervals apply to each predicted accuracy. Tables S4A–S4B give age contrasts and their CIs. Subtracting these prediction-CI endpoints does not produce a CI for the age difference.

**Supplementary Table S10C**

*Pointwise accuracy predictions at ages 50 and 85 by stage*

| Stage | Age | Predicted accuracy (%) | 95% CI |
|---|---|---|---|
| Learning | 50 | 95.39 | [94.26, 96.31] |
| Learning | 85 | 87.50 | [84.40, 90.06] |
| Novel | 50 | 75.70 | [71.57, 79.40] |
| Novel | 85 | 53.91 | [47.94, 59.77] |

Note. Predicted age-85-minus-age-50 differences are −7.89 percentage points for Learning and −21.79 for Novel. The displayed CIs concern individual predicted accuracies; Table S10D reports contrast CIs.

**Supplementary Table S10D**

*Overall and stage-specific age contrasts*

| Contrast | Measure | Estimate | 95% CI | z | p |
|---|---|---|---|---|---|
| Age 85 vs 50: overall accuracy | OR | 0.370 | [0.260, 0.526] | −5.538 | < .001 |
| Age 85 − 50: overall accuracy | Difference (pp) | −14.5 | [−19.9, −9.0] | −5.192 | < .001 |
| Age 85 − 50: Learning | Difference (pp) | −7.89 | [−11.1, −4.63] | −4.748 | < .001 |
| Age 85 − 50: Novel | Difference (pp) | −21.79 | [−29.6, −13.93] | −5.436 | < .001 |
| Novel − Learning: age differences | Difference of differences (pp) | −13.9 | [−19.4, −8.36] | −4.920 | < .001 |
| Novel / Learning: age odds ratios | Ratio of ORs | 1.11 | [0.886, 1.39] | 0.907 | .365 |
| Age 85 vs 50: correct RT | Geometric-mean ratio | 1.43 | [1.19, 1.71] | 3.864 | < .001 |

Note. Accuracy differences are in percentage points (pp); RT denotes reaction time. Intervals are pointwise and p values are unadjusted. Overall predictions average over gender, education, stage, category, and difficulty; stage-specific predictions average over the remaining factors. The stage difference of age differences and ratio of age ORs assess interaction on different scales. RT age-contrast inference is asymptotic.

**Supplementary Table S11**

*Inputs to exploratory perceptual-distinctiveness analyses*

| **Category** | **Mean SSIM** | **1 − SSIM** | **DreamSim distance** | **Age difference (pp)** |
|---|---|---|---|---|
| Animals | 0.656011 | 0.343989 | 0.462 | 8.45 |
| Bodies | 0.813484 | 0.186516 | 0.169 | 17.78 |
| Cars | 0.608356 | 0.391644 | 0.237 | 17.84 |
| Faces | 0.715313 | 0.284687 | 0.251 | 14.52 |
| Food | 0.460863 | 0.539137 | 0.508 | 9.27 |
| Hands | 0.693485 | 0.306515 | 0.264 | 20.45 |
| Houses | 0.358500 | 0.641500 | 0.338 | 6.68 |
| Musical Instruments | 0.799776 | 0.200224 | 0.508 | 17.60 |
| Tools | 0.860450 | 0.139550 | 0.380 | 9.99 |

Note. Age difference is predicted age-50 minus age-85 accuracy, using the same model estimates as Table S4. Greater positive values indicate larger later-life differences. Higher 1 − SSIM and DreamSim distance indicate greater distinctiveness. Pseudowords are excluded. Analyses use the available predictor precision and unrounded age estimates.

**Supplementary Table S12A**

*Perceptual-distinctiveness regression coefficients*

| Model / term | b | SE | 95% CI for b | p |
|---|---|---|---|---|
| **SSIM model** | | | | |
| Intercept | 13.621 | 1.542 | [9.976, 17.266] | < .001 |
| Standardized predictor | −2.639 | 1.635 | [−6.506, 1.227] | .150 |
| **DreamSim model** | | | | |
| Intercept | 13.621 | 1.559 | [9.935, 17.306] | < .001 |
| Standardized predictor | −2.560 | 1.653 | [−6.469, 1.349] | .165 |
| **Joint model** | | | | |
| Intercept | 13.621 | 1.453 | [10.066, 17.176] | < .001 |
| Standardized 1 − SSIM | −2.248 | 1.567 | [−6.082, 1.587] | .201 |
| Standardized DreamSim | −2.150 | 1.567 | [−5.985, 1.684] | .219 |

Note. *N* = 9 categories. Slopes represent percentage points per one sample-SD increase in the predictor. CIs use t distributions with 7 residual df in single-predictor models and 6 in the joint model. Intercepts describe the predicted difference at mean predictor values. All p values are unadjusted.

**Supplementary Table S12B**

*Perceptual-distinctiveness model fit and prediction error*

| Model | $R^2$ | Adjusted $R^2$ | Residual SD | AIC | BIC | LOOCV RMSE |
|---|---|---|---|---|---|---|
| SSIM | 0.271 | 0.167 | 4.625 | 56.844 | 57.436 | 5.174 |
| DreamSim | 0.255 | 0.149 | 4.676 | 57.042 | 57.634 | 5.282 |
| Joint | 0.445 | 0.260 | 4.358 | 56.389 | 57.178 | 5.531 |

Note. All models use the same nine categories. Residual SD divides residual sums of squares by residual degrees of freedom. Leave-one-category-out (LOOCV) RMSE summarizes held-out prediction errors over nine categories. Both error measures are in percentage points. AIC and BIC count residual variance as an estimated parameter. The joint-model test is $F(2, 6) = 2.41$, $p = .171$.

**Supplementary Table S13**

*Administration-mode and MoCA-completion comparisons*

| Comparison | Outcome | Estimate | 95% CI | Test | p |
|---|---|---|---|---|---|
| Online − on-site | Age (years) | −3.27 | [−8.41, 1.87] | $t(11.22) = -1.40$ | .190 |
| Online − on-site | Accuracy (pp) | 0.96 | [−2.97, 4.89] | $t(11.46) = 0.53$ | .604 |
| Online − on-site | Reaction time (s) | −0.78 | [−2.07, 0.51] | $t(9.30) = -1.36$ | .205 |
| MoCA completion | Age (years) | 0.12 | [−3.36, 3.61] | $t(86.57) = 0.07$ | .943 |

| **Comparison** | **Outcome** | **Estimate** | **95% CI** | **Test** | **p** |
|---|---|---|---|---|---|
| MoCA completion | Accuracy (pp) | −0.38 | [−3.10, 2.33] | $t(90.81) = -0.28$ | .780 |
| MoCA completion | Reaction time (s) | 0.01 | [−0.59, 0.62] | $t(88.69) = 0.04$ | .971 |
| MoCA completion | Male vs female | OR 1.083 | [0.467, 2.513] | Fisher exact | 1.000 |
| MoCA completion | Higher vs lower education | OR 1.773 | [0.702, 4.479] | $\chi^2(1) = 0.96$ | .326 |

Note. Continuous estimates are raw mean differences with Welch CIs. Mode differences are online minus on-site; MoCA differences are completers minus non-completers. Categorical estimates are unadjusted odds ratios from counts given in S3.4, with log-Wald 95% CIs; tests retain the original Fisher exact or $\chi^2$ p values. The small samples and nonsignificant tests do not establish equivalence.

**Supplementary Table S14**

*Pairwise orientation contrasts within stage*

| Stage | Contrast | OR | Bonferroni 95% CI | z | Holm p |
|---|---|---|---|---|---|
| Learning | Frontal / Left30 | 0.711 | [0.647, 0.780] | −8.76 | < .001 |
| Learning | Frontal / Right30 | 0.723 | [0.657, 0.794] | −8.24 | < .001 |
| Learning | Left30 / Right30 | 1.017 | [0.919, 1.125] | 0.40 | .691 |
| Novel | FrontalS1 / FrontalS2 | 1.050 | [0.972, 1.120] | 1.71 | .088 |
| Novel | FrontalS1 / Left60 | 1.280 | [1.195, 1.380] | 9.84 | < .001 |
| Novel | FrontalS1 / Right60 | 1.370 | [1.272, 1.470] | 11.97 | < .001 |
| Novel | FrontalS1 / RightS1 | 1.490 | [1.388, 1.600] | 15.53 | < .001 |
| Novel | FrontalS2 / Left60 | 1.230 | [1.145, 1.320] | 8.19 | < .001 |
| Novel | FrontalS2 / Right60 | 1.310 | [1.218, 1.410] | 10.33 | < .001 |
| Novel | FrontalS2 / RightS1 | 1.430 | [1.329, 1.530] | 13.88 | < .001 |
| Novel | Left60 / Right60 | 1.070 | [0.995, 1.140] | 2.62 | .017 |
| Novel | Left60 / RightS1 | 1.160 | [1.085, 1.250] | 6.10 | < .001 |
| Novel | Right60 / RightS1 | 1.090 | [1.016, 1.170] | 3.42 | .002 |

Note. OR compares the first condition with the second. Familywise 95% CIs use Bonferroni adjustment within stage (three Learning and ten Novel contrasts); p values use Holm adjustment within those same families. Different adjustments explain why the Left60 / Right60 CI includes 1 although its Holm p value is .017. Condition definitions are in Table S10A.
